# Small RNA Sequencing Reveals a Distinct MicroRNA Signature between Glucocorticoid Responder and Glucocorticoid Non-responder Primary Human Trabecular Meshwork Cells after Dexamethasone Treatment

**DOI:** 10.1101/2023.06.19.545545

**Authors:** Kandasamy Kathirvel, Xiaochen Fan, Ravinarayanan Haribalaganesh, Devarajan Bharanidharan, Rajendrababu Sharmila, Ramasamy Krishnadas, Veerappan Muthukkaruppan, Colin E. Willoughby, Srinivasan Senthilkumari

**Affiliations:** Department of Ocular Pharmacology, Aravind Medical Research Foundation, Madurai, Tamilnadu, India; Institute of Life Course and Medical Sciences, University of Liverpool, England, United Kingdom; Department of Bioinformatics, Aravind Medical Research Foundation, Madurai, Tamilnadu, India; Glaucoma Clinic, Aravind Eye Hospital, Madurai, Tamilnadu, India; Advisor, Aravind Medical Research Foundation, Madurai, Tamilnadu, India; Genomic Medicine, Biomedical Sciences Research Institute, Ulster University, Northern Ireland, United Kingdom

## Abstract

The present study aimed to understand the role of miRNAs in differential glucocorticoid (GC) responsiveness in human trabecular meshwork (HTM) cells using small RNA sequencing. For this, total RNA was extracted from cultured HTM cells with known GC responsiveness using Human organ-cultured anterior segment (HOCAS) (GC-responder GC-R; n=4) and GC-non-responder (GC-NR; n=4) after treatment with either 100nM dexamethasone (DEX) or ethanol (ETH) for 7 days. Differentially expressed miRNAs (DEMIRs) were compared among 5 groups and validated by RT-PCR. There were 13 and 21 DEMIRs identified in Group #1 (ETH vs DEX-treated GC-R) and Group #2 (ETH vs DEX-treated GC-NR) respectively. Seven miRNAs were found as common miRNAs dysregulated in both GC-R and GC-NR (Group #3). There were 6 and 14 unique DEMIRs were identified in GC-R (Gropu#4) and GC-NR (Group#5) HTM cells respectively. Ingenuity Pathway Analysis identified enriched pathways and biological processes associated with differential GC responsiveness in HTM cells. Integrative analysis of miRNA-mRNA of the same set of HTM cells revealed several molecular regulators for GC non-responsiveness. This is the first study revealed a unique miRNA signature between GC-R and GC-NR HTM cells which raises the possibility of developing new molecular targets for the management of steroid-OHT/glaucoma.

## Introduction

Glucocorticoids (GCs) are known to regulate several physiological processes and are the mainstay in the management of inflammatory eye diseases (Gordon, 1955). Long-term use of GC causes raised intraocular pressure (IOP) or ocular hypertension (OHT) in about 30-50% of the susceptible individuals depending on the route of administration. Steroid-induced OHT if left un-treated can lead to secondary open angle glaucoma (Becker and Mills, 1963). Individuals with GC sensitivity are more likely to develop the most prevalent form of glaucoma i.e. primary open angle glaucoma (POAG); affected people show more than 90% susceptiblility to steroid-induced raised IOP (Bartlett et al., 1993; Lewis et al., 1988). Even though POAG and steroid-induced glaucoma (SIG) share similar clinical presentations, the molecular mechanism responsible for the differential GC responsiveness in individuals is not well understood.

MicroRNAs (miRNAs) are small non-coding RNAs which regulate gene expression by either mRNA degradation or translational repression(Guo et al., 2010). MiRNAs have been detected in most of the biological fluids where they are preserved in micro-vesicles, exosomes or bound to carrier proteins thereby providing a remarkable stability to miRNAs (Mitchell et al., 2008). These properties makes miRNAs suitable bio-markers for many diseases including ocular diseases (Kumar and Reddy, 2018; Liu et al., 2018; Mitchell et al., 2008; Kim and Zhang, 2019).

In the eye, miRNAs are expressed in tissue-specific fashion and have a specific role in ocular development and retinal homeostasis (Hackler et al., 2010; Busskamp et al., 2014). MiRNAs have been identified in ocular fluids such as tears, aqueous humor and vitreous humor (Liu et al., 2018). Several studies documented the expression of glaucoma-associated miRNAs in affected tissues such as aqueous humor, tears, trabecular meshwork (TM) and retina of patients with glaucoma and animal models (Liu et al., 2018; Tanaka et al., 2014; Jayaram et al., 2017; Hubens et al., 2021; Kosior-Jarecka et al., 2021; Greene et al., 2022). The miRNA expression profile of the TM in response to patho-physiologically relevant stressors such as cyclic mechanical stress, oxidative stress and stress-induced premature senescence has been reported and contribute to vital cellular functions (Polansky et al., 1991; Li et al., 2010, 2009; Zhao et al., 2019; Shen et al., 2020; Youngblood et al., 2020).

GC production by the adrenal glands and GC-mediated cellular response are also regulated by miRNAs (Clayton et al., 2018). In the TM, dexamethasone (DEX) treatment induces cellular and extracellular remodeling leading to increased outflow resistance and elevated IOP (Yemanyi et al., 2020). The pharmacological actions of GCs are mediated by glucocorticoid receptor whose activation results in the stimulation of target gene expression that regulates several complex signaling pathways (Whirledge and DeFranco, 2018; Kathirvel et al., 2022). The contribution of miRNA in the regulation of GC activity and signaling that results in the differential GC responsiveness in the TM is currently unknown. Therefore, in this study the differential expression of miRNAs in primary HTM cells with known GC responsiveness was investigated using small RNA sequencing. A unique miRNA signature between GC-R and GC-NR HTM cells was identified and understanding the role of miRNAs in GC responsiveness raises the possibility of new molecular targets for the management of steroid-induced OHT/glaucoma.

## Methods

### Human Donor Eyes

Post-mortem human cadaveric eyes not suitable for corneal transplantation were obtained from the Rotary Aravind International Eye Bank, Aravind Eye Hospital, Madurai. The study was conducted following approval from the Human Ethics Committee of the Institute. The tissues were handled in accordance with the Declaration of Helsinki. The donor eyes were enucleated within 5 h of death (mean elapsed time between death and enucleation was 2.75 ±1.58 h) and kept at 4° C in the moist chamber until culture. All eyes were examined under the dissecting microscope for any gross ocular pathological changes and only macroscopically normal eyes were used for the experiments. The characteristics of donor eyes used for this study is summarized in Supplementary Table S1.

### Primary Human Trabecular Meshwork (HTM) Cells with Known GC responsiveness

In a set of paired eyes, one eye was used to establish HOCAS *ex vivo* model system to characterize GC responsiveness after DEX treatment as prescribed previously (Haribalaganesh et al., 2021). The other eye was used to establish primary HTM cultures from eyes with identified GC responsiveness (Kathirvel et al., 2022). The TM tissue was excised from the other eye of each set of paired eyes and primary HTM cell culture was established by extracellular matrix digestion method as described previously (Stamer et al., 1995; Ashwinbalaji et al., 2018).

Primary HTM cells were grown at 37° C in 5% CO_2_ in low glucose Dulbecco’s modified Eagle medium (DMEM) with 15% fetal bovine serum, 5 ng/ml basic fibroblast growth factor and antibiotics. The primary HTM cells isolated from the other eye of each pair were characterized with aquaporin, myocilin and phalloidin staining by immunofluorescence analysis (data not shown). HTM cells with more than 50% myocilin positivity in response to DEX treatment were used for further experiments (Keller et al., 2018). Confluent cultures of GC-R and GC-NR HTM cells were then treated with either 100nM DEX or 0.1% ethanol (ETH) as a vehicle control over a period of 7 days and the medium was exchanged every other day (3 doses of DEX treatments). HTM cells from passages 2-4 were used for all experiments. At the end of DEX or 0.1% ETH treatment for 7 days, HTM cells from each GC-R (n=4) and GC-NR (n=4) HTM cells were subjected to RNA extraction and small RNA sequencing.

### Total RNA extraction

Total RNA including miRNA was extracted from GC-R and GC-NR HTM cells post 7d DEX treatment using the TRIzol reagent. Briefly, 200µl of chloroform was added to 1ml of TRIzol containing one million HTM cells. Then, 500µl of isoproponal and 1µl of glycogen were added into RNA containing aqueous phase for nucleic acid precipitation after centrifugation at 13000rpm for 15mins at 4°C. RNA pellet was washed twice with 75% ethanol and eluted with 20µl of nucleus free water. RNA quantity and quality were assessed by the ratio of absorbance at 260/280 nm using NanoDrop 2000 spectrophotometer (Thermofisher Scientific, DE, UK), and TapeStation (Agilent Technologies, CA, USA), respectively.

### Library preparation and miRNA sequencing

MiRNA sequencing libraries were prepared using NEBNext smallRNA library prep kit (NEB, MA, USA). Briefly, RNA fragments of different sizes were separated by PAGE and 18 to 30 nt stripe was ligated with specific adapters at 3’ and 5’ end. Then, libraries were reverse transcribed followed by PCR amplification. The final enriched libraries were purified and quantified by Qubit and the size was analyzed by Bio analyzer. Individual libraries were pooled with specific index sequences and then sequenced by Illumina Next Seq 500 platform (Illumina, Inc., San Diego, CA, USA), according to the manufacturers protocol. Around 10 million reads were generated from each HTM donor RNA sample and obtained through de-multiplexing.

### Differential Expression Analysis of miRNAs

The quality of raw data of miRNA sequencing was assessed by FastQC tool. The PCR duplicates and low-quality reads were excluded using cutadapt3 in the data cleaning step. The pre-processed high quality miRNA reads were then mapped with human reference genome GRCh38 using STAR aligner with default parameters. The expression of miRNAs in read count was obtained from individual aligned BAM files using FeatureCounts with miRbase annotation (release 22). MiRNAs with less than five read count were excluded for further analysis. TMM (timed mean of M values) strategy was employed for data normalization and differential expression was performed using EdgeR, package from Bioconductor. The miRNAs were considered as differentially expressed if the absolute fold change (log2) value was more than 1.5, and the P value < 0.05. For comparison, the differentially expressed miRNAs were segregated into 5 groups as described previously (Kathirvel et al., 2022). Group #1 = DEMirs between DEX and ETH treated GC-R HTM cells; Group #2 = DEMirs between DEX and ETH treated GC-NR HTM cells; Group #3 = Commonly expressed miRNAs between Group #1 and Group #2; Group #4 = Uniquely expressed miRNAs in GC-R HTM cells (Group #4 = Group #3 minus Group #1); Group #5 = Uniquely expressed miRNAs in GC-NR HTM cells (Group #5 = Group #3 minus Group #2).

### Validation of DEMirs by Real Time-PCR

The expression of significantly altered miRNAs identified by small RNA sequencing analysis was further validated by RT-qPCR with additional, independent biological replicates of GC-R and GC-NR (n=5 each) HTM cells. Briefly, the total RNA was extracted as described previously (Kathirvel et al., 2022) and miRNAs from each HTM cells was reverse transcribed into cDNA using miScript II RT kit as per the manufacturer’s instruction (Qiagen, Hilden, Germany). RT-qPCR was performed in a total volume of 20 µl containing 25 ng of total cDNA, 5x SYBR Green master mix and final concentration of primers. The assay was performed on an ABI-QuantStudio 5 (Applied Biosystems, MA, USA). All miRNA levels were measured at CT threshold levels and normalized with the average CT values of reference target RNU6. Values were expressed as fold increase over the corresponding values for control by the 2^-ΔΔCT^ method. All PCR reactions were performed in triplicates.

### Prediction of the target mRNA with their enriched pathways and biological processes

The target mRNAs of DEMirs were predicted using Ingenuity Pathway Analysis software (IPA, Qiagen, UK). The DEMirs from all groups (Group #1 to #5) with P-value<0.05 were analyzed using the ‘miRNA Target Filter’ tool in the IPA software based on the TargetScan, TarBase, miRecords contents and Ingenuity® Knowledge Base. The cumulative weighted context score (CWCS) defined by TargetScan was used for assigning different confidence levels of the predicted target mRNAs. The target mRNAs with ‘experimentally observed’ and ‘high confidence level’ were selected into the ‘Target MRNA List 1’.

At the time of miRNA extraction using the same experimental set-up mRNA was also extracted. This provided paired mRNA-seq and miRNA-seq data from the same cells of known GC responsiveness. The mRNA-seq was previously published by our group (Kathirvel et al., 2022). The DEMirs from Group #1 to #5 were further therefore combined with our previously generated mRNA library (Kathirvel et al., 2022) to find the negatively correlated miRNA targets (Target MRNA List 2) using the IPA software in paired datasets from the same TM cell line donors. The prediction of the pathways and biological processes enriched in the ‘Target MRNA List 1’ and ‘Target MRNA List 2’ was performed using the GSEApy package /python (v 3.8.3) (for ‘Target MRNA List 1’ without logFC and p-values) (Fang et al., 2023) and/or IPA software (for ‘Target MRNA List 2’ with logFC and p-values).

### Statistical Analysis

Statistical analysis was carried out using Graph Pad Prism (ver.8.0.2) (Graph Pad software, CA, USA). All data are presented as mean ± SEM or otherwise specified. Statistical significance between two groups was analyzed using unpaired 2-tailed Student’s t test. P <0.05 or less was considered as statistically significant.

## RESULTS

### Establishment of GC-R and GC-NR HTM cells

In this study, one eye of each set of paired eyes was used to assess the GC responsiveness in HOCAS after 100nM DEX treatment for 7 days and the other eye was used to establish the primary HTM cells. Based on the IOP response, the HTM cells established from each donor eye was categorized as GC-R or GC-NR cells (Haribalaganesh et al., 2021; Kathirvel et al., 2022). The details of human donor eyes used in this study are summarized in Supplementary Table S1.

### miRNA Seq Data Quality

In total, 9.1 to 17.2 million reads were generated from miRNA sequencing for each primary HTM cells. Pre-aligned QC reports showed the quality score (Phred or Q score) of miRNA reads were ≥ 30. An average of 97% reads from miRNA-seq data were aligned with human reference genome GRCh38. The details of miRNA-sequencing and alignment statistics are shown in Supplementary Table S2. EdgeR with Benjamini-Hochberg Corrections was used to identify the differentially expressed miRNAs from all five groups (Table 1; Supplementary Table S3a&S3b).

**Table 1.**
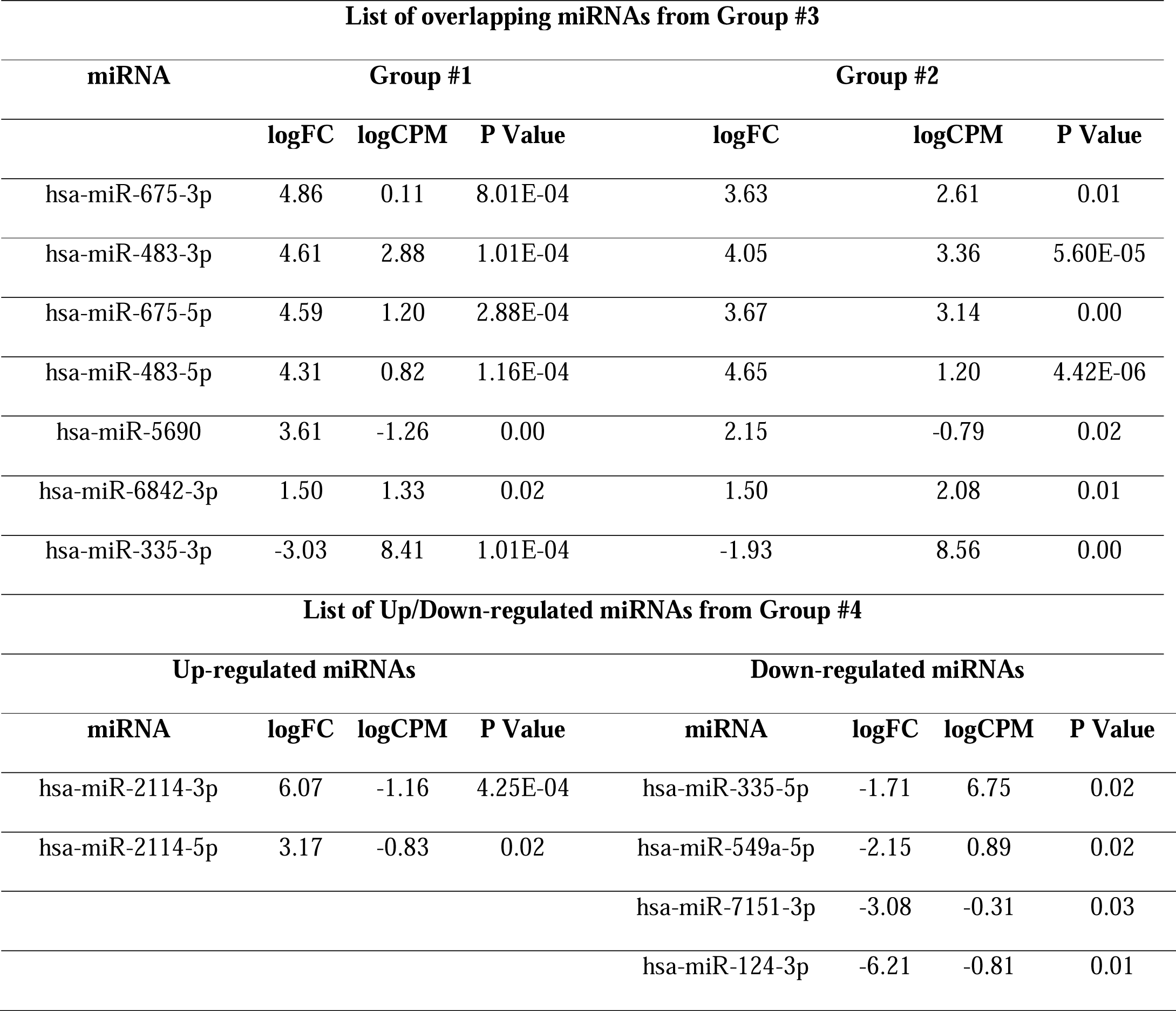
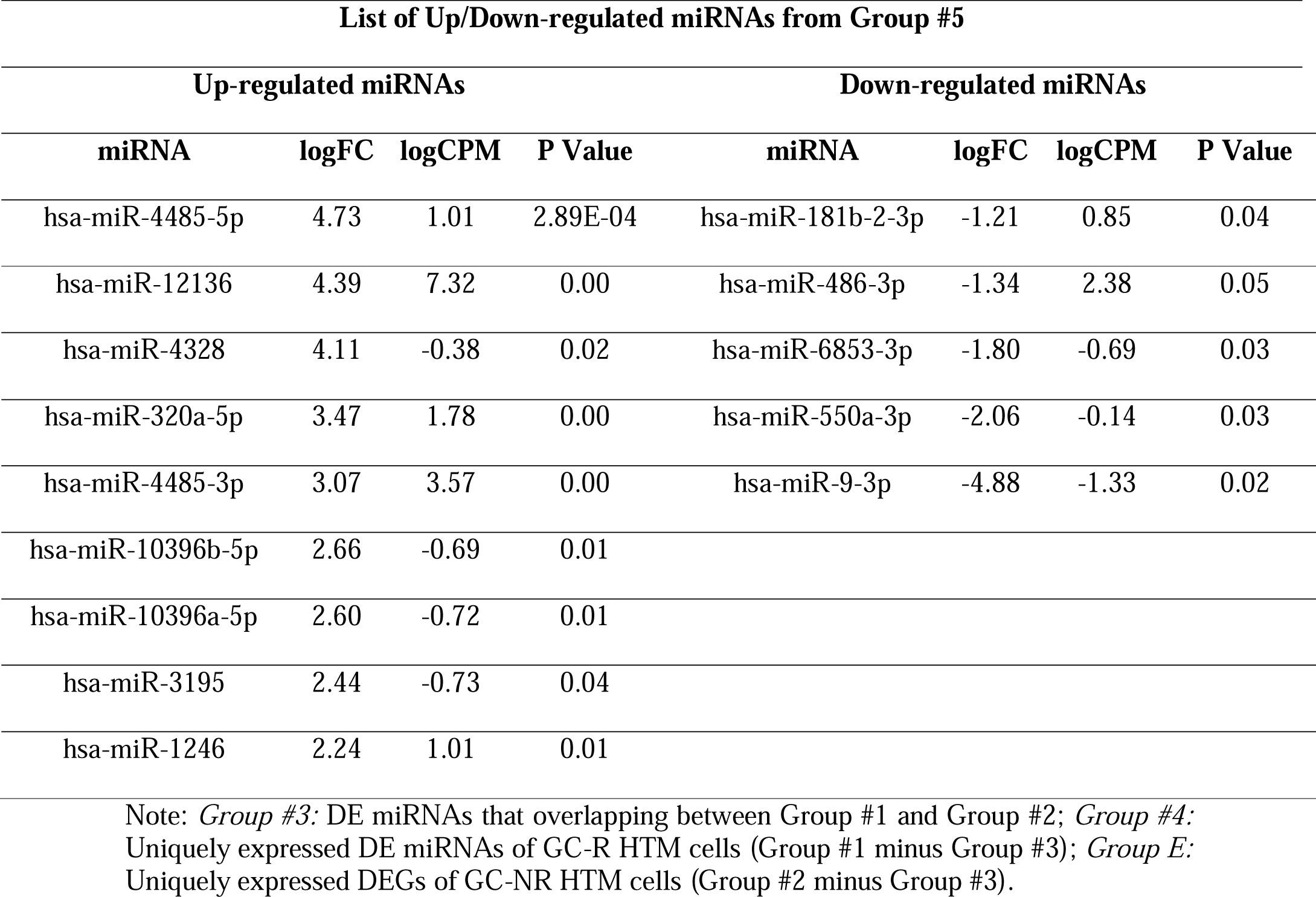
List of overlapping and Up/Down-regulated miRNAs from Group #3, #4 and #5.

### Differentially expressed genes of GC-R and GC-NR HTM cells

The total number of miRNAs identified in HTM cells of each donor eye ranged from 718 to 898. The expression of DE-miRNAs from GC-R (Group #1) and GC-NR (Group #2) HTM cells are represented in volcano plot (Figure 1). In total, there were 13 and 21 miRNAs identified as differentially expressed in Group #1 (8 up-regulated; 5 down-regulated) and Group #2 (15 up-regulated; 6 down-regulated), respectively. Seven miRNAs were found as common miRNAs dysregulated in both GC-R and GC-NR (Group #3) with absolute fold change (log2) value 1.5, and the *P value* < 0.05. In total, 6 (2 up-regulated; 4 down-regulated) and 14 (9 up-regulated; 5 down-regulated) miRNAs were found to be uniquely expressed only in GC-R (Group #4) and GC-NR (Group #5) HTM cells, respectively (Figure 2). In Group #4 (GC-R), hsa-miR-2114-3p (log FC=6.07), hsa-miR-2114-5p (log FC=3.17) were significantly up-regulated and hsa-miR-335-5p (log FC= -1.71), hsa-miR-549a-5p (logFC= - 2.15), hsa-miR-7151-3p (logFC=-3.08) and hsa-miR-124-3p (logFC=-6.21) were significantly down-regulated. Whereas in Group #5 (GC-NR), hsa-miR-4485-5p (logFC=4.73), hsa-miR-12136 (logFC=4.39), hsa-miR-4328 (logFC=4.11) were significantly up-regulated and hsa-miR-181b-2-3p (logFC= -1.21), hsa-miR-486-3p (logFC=-1.34) and hsa-miR-6853-3p (logFC=-1.80) were significantly down-regulated. In Group #3, the commonly expressed miRNAs between Group #1 and #2 were hsa-miR-675-3p, hsa-miR-483-3p, hsa-miR-675-5p, hsa-miR-483-5p, hsa-miR-5690, hsa-miR-6842-3p (up-regulated); and hsa-miR-335-3p (down-regulated). The DEMirs from Group #1 to Group #5 are shown in Venn diagram (Figure 2) and Table 1 and Supplementary Table S3a& S3b.

**Figure 1.**
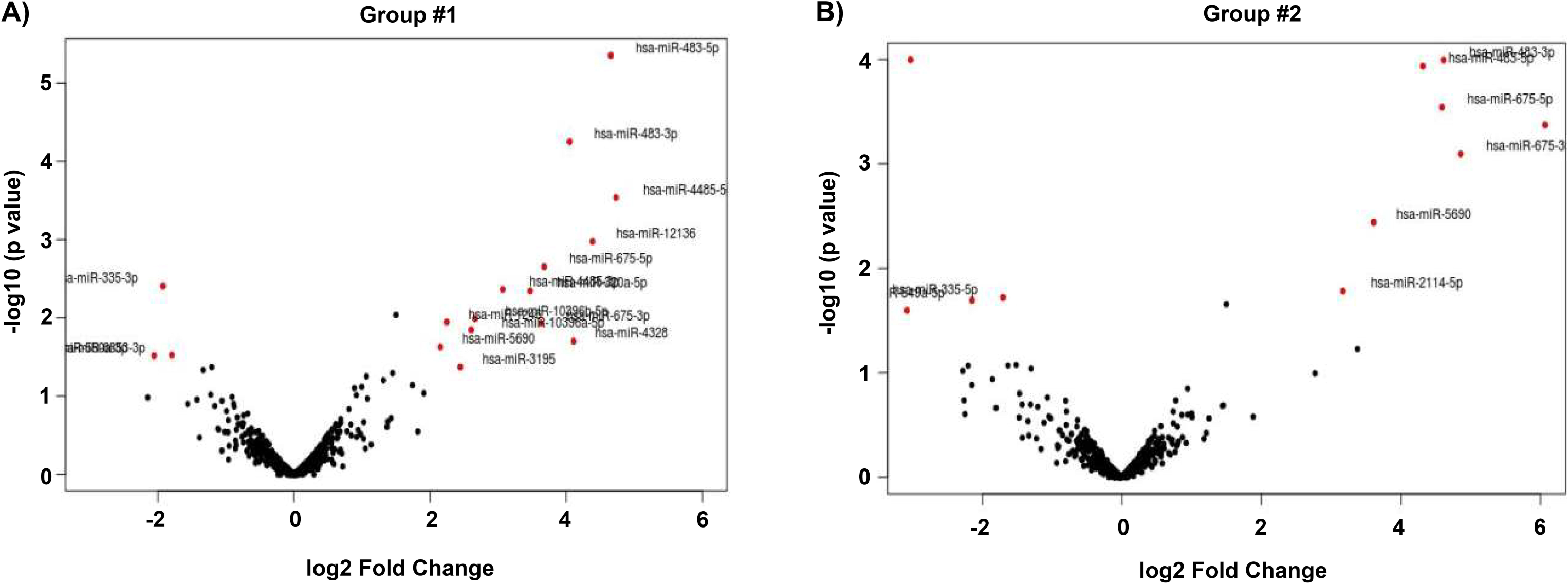
Volcano plot showing the Distribution of DE miRNAs. The fold of change (log2) and p value (-log 10) of the dysregulated miRNAs in DEX treated cells compared to vehicle control are shown in volcano plot. Significant p value <0.05, only taken for consideration (red color).

**Figure 2.**
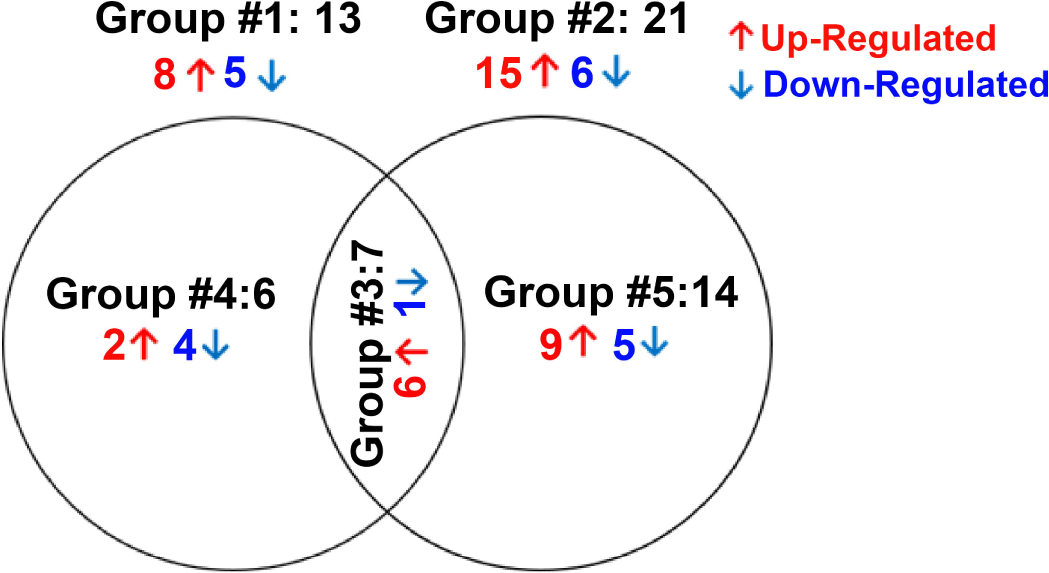
Venn diagram Showing Differentially Expression Groupings. DE miRNAs of three groups from miRNA seq data are shown. Only genes with absolute fold change 1.5 and significant *p* value <0.05 were included in these groupings. Group #1: DE miRNAs between ETH and DEX-treated cells of GC-R HTM cells, Group #2: DE miRNAs between ETH and DEX-treated cells of GC-NR HTM cells, Group #3: Overlapping DE miRNAs between Group #1 and Group #2; Group #4: uniquely expressed miRNAs in GC-R and Group #5: uniquely expressed DE miRNAs in GC-NR.

#### Validation of DE-miRNAs by qPCR

Out of 9 miRNAs selected for qPCR (Supplementary Table S5), the expression pattern of 7 miRNAs matched with miRNA seq data (Figure 3; Figure 4).

**Figure 3.**
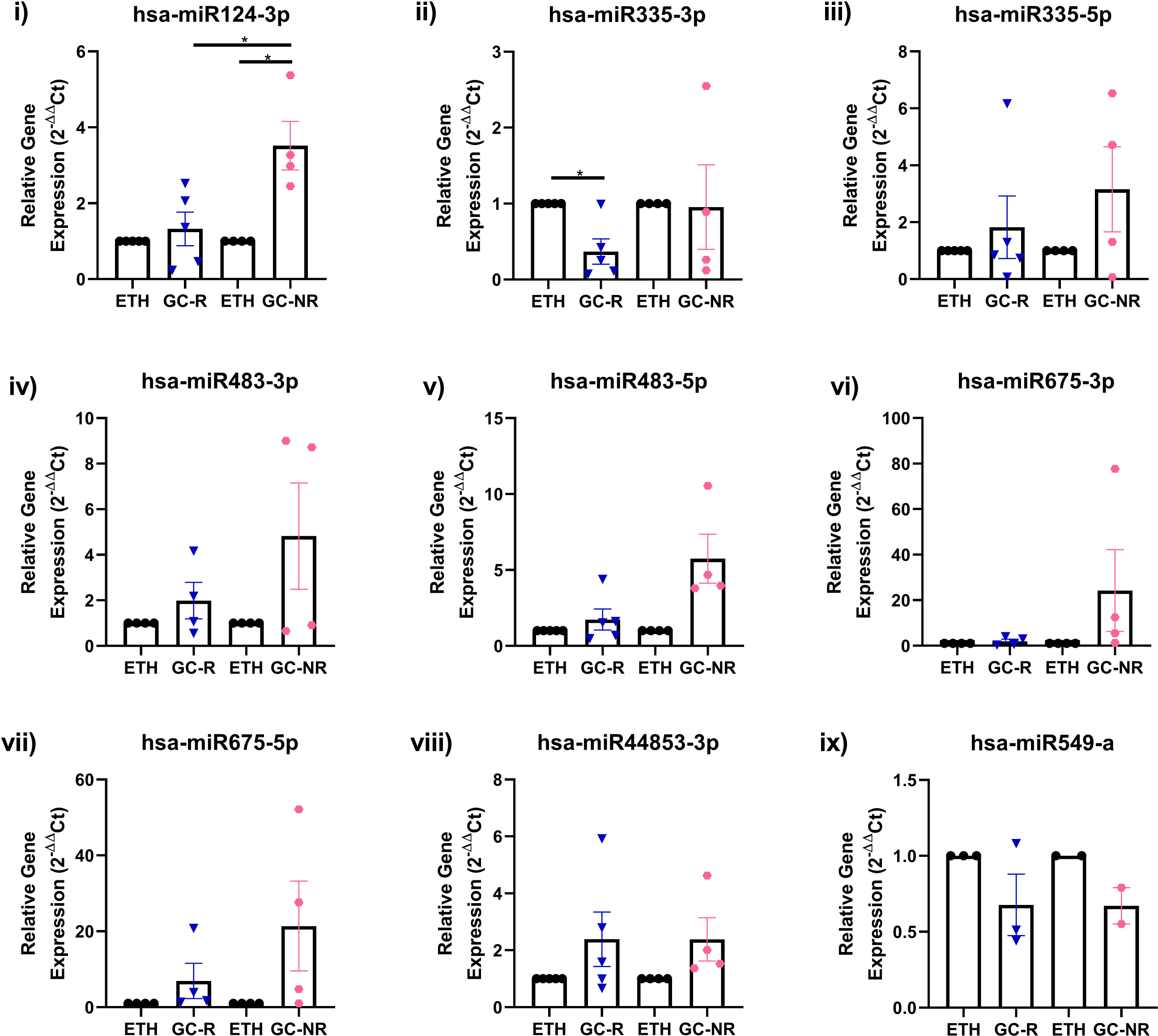
Validation of DE miRNAs by qPCR. Expression profile of selected miRNAs identified from miRNA-seq was validated by qPCR is shown. Primary HTM cells were treated with 100nM DEX or 0.1% ETH for 7 days. Total RNA was extracted, converted to cDNA and the expression profile of selected miRNAs were carried out qPCR. MiRNA expressions were normalized to RNU6, and analyzed using the 2^−ΔΔCT^ method.

**Figure 4.**
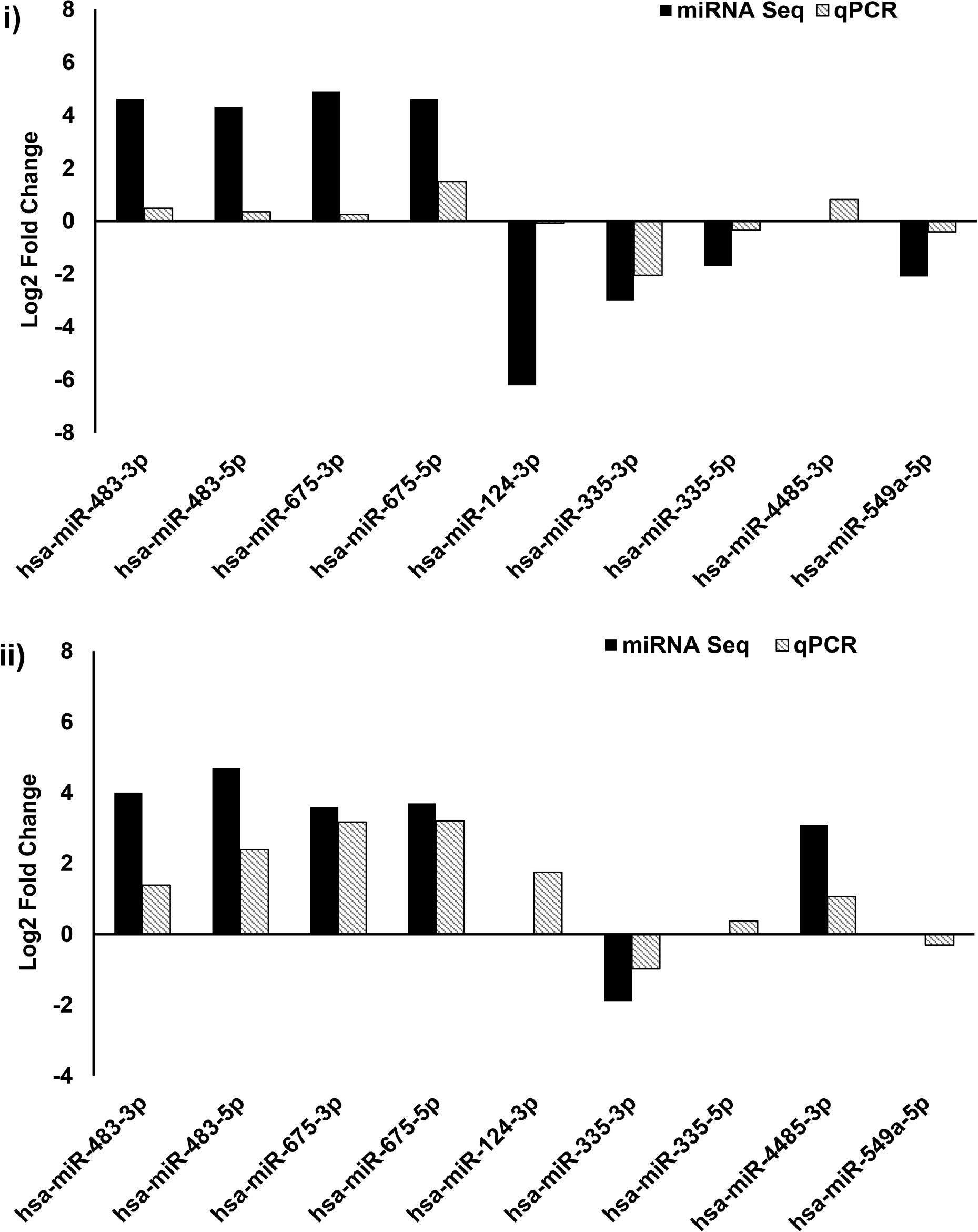
Validation of miRNA sequencing by qPCR. Comparison of selected genes expression profile from Group #1 (i) and Group #2 (ii) miRNA sequence by qPCR; 2^ΔΔCt^ method was used for calculation of miRNA expression changes and U6 was used as an internal control.

### Prediction of target mRNAs and Pathways Analysis

#### A) Prediction of target mRNAs and pathways in silico analysis-‘Target MRNA List 1’

The prediction of the target mRNAs in the ‘Target MRNA List 1’ (without logFC and p-values) was performed by the IPA software based on the TargetScan, TarBase, miRecords contents and Ingenuity® Knowledge Base. We found 1354 target mRNAs (74 of 1354 miRNA: mRNA interactions were experimentally validated) in Group #3, 1173 target mRNAs (237 of 1173 miRNA: mRNA interactions were experimentally validated) in Group #4, and 3187 target mRNAs (5 of 3187 miRNA: mRNA interactions were experimentally validated) in Group #5 (Supplementary File 1_List 1A). The enriched pathways and biological processes in the ‘Target MRNA List 1’ were predicted using the GSEApy package

/python (v 3.8.3) (Fang et al., 2023) based on the KEGG_2021 and GO_Biological_Process_2021 datasets. We found 27 significant pathways and 266 biological processes in Group #3, 46 significant pathways and 360 biological processes in Group #4, and 47 significant pathways and 445 biological processes in Group #5 (p-value<0.05) (Supplementary file1_List 1B and 1C). The top 10 KEGG pathways and biological processes in Group #4-#5 based on the smallest p-value are shown in Figure 5.

**Figure 5.**
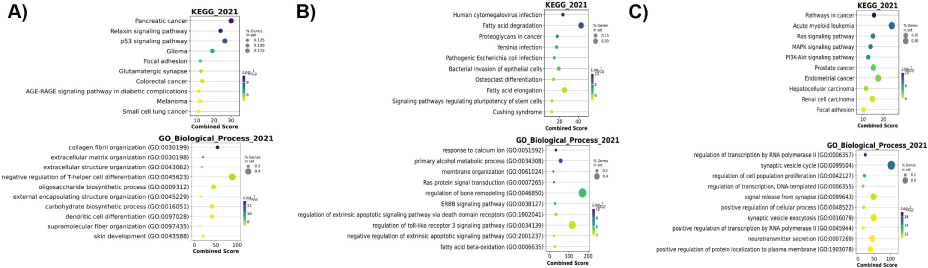
Top 10 predicted KEGG pathways and GO biological processes of the ‘Target MRNA List 1’ of Group #3 (A), #4 (B) and #5 (C). Dot colours: Log (1/P-value). Dot sizes: % genes in set. The combined score is defined based on the percentages of genes in set and Log (1/P-value) by [3]. P-value<0.05 was considered to be statistically significant.

#### B) Prediction of target mRNAs that negatively correlated with DEMirs, their pathways and biological processes in experimental analysis – ‘Target MRNA List 2’

The target mRNAs that negatively correlated with the DEMIRs ‘Target MRNA List 2’ (with logFC and p-values) were predicted by the IPA software based on our previously generated mRNA library (Kathirvel et al., 2022). We found 2 target mRNAs were negatively correlated with 2 DEMirs in Group #3, 15 target mRNAs were negatively correlated with 4 DEMirs in Group #4, and 12 target mRNAs were negatively correlated with 6 DEMirs in Group #5 (Table 2). The confidence levels of each predicted target mRNAs are shown in Supplementary file 2_List 2A. The interaction networks of the DEMirs from Group #4 and #5 and their negatively correlated mRNAs were shown in the Figure 6. The pathway prediction of the ‘Target MRNA List 2’ was performed using the GSEApy package /python (v 3.8.3) (Supplementary file 2_List 2B) and IPA software (Supplementary file 2_List 2C). The GSEApy pathway analysis results of ‘Target MRNA List 2’ (Supplementary file 2_List 2B) were compared to that of ‘Target MRNA List 1’ (Supplementary file1_List 1B) for finding the overlapping pathways. We found the MAPK signaling pathway was statistically significant in ‘Target MRNA List 1&2’ of Group #3 (p-value<0.05). The nicotinate and nicotinamide metabolism pathway was statistically significant both in ‘Target MRNA List 1&2’ of Group #5. No significant overlapping pathway was found in the ‘Target MRNA List 1&2’ of Group #4.

**Figure 6.**
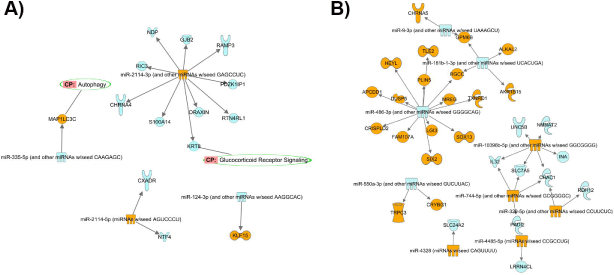
Interaction networks of the DEMIRs from Group #4 (A) and Group #5 (B) (absolute LogFC>2 and P<0.05) and their negatively corelated target mRNAs (absolute FC>2 and P<0.05). Orange: upregulated miRNAs/mRNAs. Blue: downregulated miRNAs/mRNAs. Green circle: predicted pathways that the target mRNAs are involved in.

**Table 2.** showing the target mRNAs that negatively correlated with DEMIRs in Group#3-#5. RGS16: Regulator of G Protein Signaling 16; NGFR: Nerve Growth Factor Receptor; KLF15: KLF Transcription Factor 15; CHRNA4: Cholinergic Receptor Nicotinic Alpha 4 Subunit; DRAXIN: Dorsal Inhibitory Axon Guidance Protein; GJB2: Gap Junction Protein Beta 2; JCHAIN: Joining Chain of Multimeric IgA And IgM; KRT8: Keratin 8; NDP: Norrin Cystine Knot Growth Factor NDP; PDZK1IP1: PDZK1 Interacting Protein 1; RAMP3: Receptor Activity Modifying Protein 3; RIC3: RIC3 Acetylcholine Receptor Chaperone; RTN4RL1: Reticulon 4 Receptor Like 1; S100A14: S100 Calcium Binding Protein A14; CXADR: CXADR Ig-Like Cell Adhesion Molecule; NTF4: Neurotrophin 4; MAP1LC3C: Microtubule Associated Protein 1 Light Chain 3 Gamma; CHAC1: ChaC Glutathione Specific Gamma-Glutamylcyclotransferase 1; INA: Internexin Neuronal Intermediate Filament Protein Alpha; NMNAT2: Nicotinamide Nucleotide Adenylyltransferase 2; UNC5B: Unc-5 Netrin Receptor B; ALKAL2: ALK And LTK Ligand 2; SLC24A2: Solute Carrier Family 24 Member 2; DUSP5: Dual Specificity Phosphatase 5; LGI3: Leucine Rich Repeat LGI Family Member 3; MREG: Melanoregulin; SOX13: SRY-Box Transcription Factor 13; IL32: Interleukin 32; and SLC7A5: Solute Carrier Family 7 Member 5.

| *(i) Target mRNAs that negatively correlated with DEMIRs in Group#3* |  |  |
| --- | --- | --- |
| **ID of DEMIRs** | **Count of Target mRNAs** | **ID of Target mRNAs** |
| **hsa-miR-483-5p** | 1 | RGS16 |
| **hsa-miR-6842-3p** | 1 | NGFR |
| *(ii) Target mRNAs that negatively correlated with DEMIRs in Group#4* |  |  |
| **hsa-miR-124-3p** | 1 | KLF15 |
| **hsa-miR-2114-3p** | 11 | CHRNA4, DRAXIN, GJB2, JCHAIN, KRT8, NDP, PDZK1IP1, RAMP3, RIC3, RTN4RL1, S100A14 |
| **hsa-miR-2114-5p** | 2 | CXADR, NTF4 |
| **hsa-miR-335-5p** | 1 | MAP1LC3C |
| *(i) Target mRNAs that negatively correlated with DEMIRs in Group#5* |  |  |
| **hsa-miR-10396b-5p** | 4 | CHAC1, INA, NMNAT2, UNC5B |
| **hsa-miR-181b-2-3p** | 1 | ALKAL2 |
| **hsa-miR-320a-5p** | 1 | CHAC1 |
| **hsa-miR-4328** | 1 | SLC24A2 |
| **hsa-miR-486-3p** | 4 | DUSP5, LGI3, MREG, SOX13 |
| **hsa-miR-10396a-5p** | 3 | CHAC1, IL32, SLC7A5 |

The comparative analysis of the pathways of the ‘Target MRNA List 2’ in Group #3, #4 and #5 was performed by IPA software (Supplementary file 2_List 2D). The top 20 significant pathways in Group #3, #4 and #5 ranked by -log(P-value) are shown in Figure 7A. The results show that Neurotrophin/TRK signaling was significant in Group #3 (-log(p-value) =2.18) and #4 (-log(p-value) =1.34). NAD signaling pathway (-log(p-value) =2.67), NAD biosynthesis III (-log(p-value) =2.55), NAD salvage pathway III (-log(p-value) =2.49), NAD biosynthesis from 2-amino-3-carboxymuconate semialdehyde (-log(p-value) =2.49), NAD biosynthesis II (from tryptophan) (-log(p-value) =2.22), and γ-glutamyl cycle (-log(p-value) =2.22) pathways were found to be significant only in Group #5 (Supplementary file 2_List 2D).

**Figure 7.**
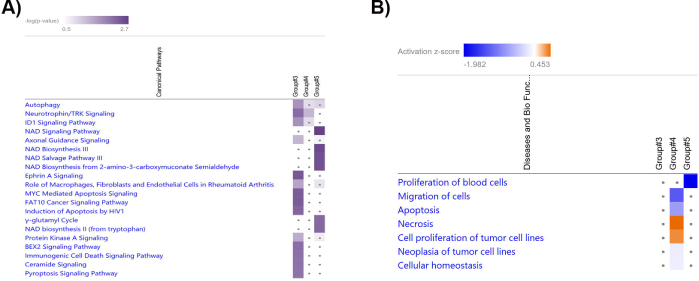
Heatmap shows the top 20 significant pathways (A) and biological processes (B) in the comparative analysis of Group #3, #4 and #5. Purple colours: -log (p-value). Darker colour: higher -log (p-value). Grey dots: pathways with p-value>0.05. Right-tailed Fisher’s Exact test was used for statistical analyses. P-value<0.05 was considered to be statistically significant. Biological processes are ranked according to the absolute z-score. Orange: activation z-score>0. Blue: activation z-score<0. White: z-score=0. Grey dots: P-value>0.05. Right-tailed Fisher’s Exact test was used for statistical analyses. P-value<0.05 was considered to be statistically significant.

The biological process prediction of the ‘Target MRNA List 2’ was performed using the GSEApy package /python (v 3.8.3) and IPA software (Supplementary file 2_List 2E). The comparative analyses of the GSEApy biological processes of the ‘Target MRNA List 1 & 2’ were performed using python (v 3.8.3). The overlapping biological processes in the ‘Target MRNA List 1 & 2’ of Group #3-#5 are shown in Supplementary Table S6.

The biological processes of the ‘Target MRNA List 2’ were further analyzed and compared by IPA software. The comparative analysis of the biological processes in Group #3, #4, and #5 are shown in Figure 7 (B) (ranked by absolute z-score. z-score is to ascertain the activation states of involved biological functions; z>0: increased and z<0: decreased)). The results show that the cell migration (Z-score= -1.295), apoptosis (Z-score= -0.765), and cellular homeostasis (Z-score= -0.147) related biological processes were predicted to be down-regulated in Group #4, while necrosis related biological process (Z-score=0.453) were up-regulated in Group #4 compared to the Group #3 and #5.

## Discussion

Steroid-induced OHT and glaucoma are serious consequences associated with the long-term use of steroids with many ocular conditions requiring chronic steroid administration (Fini et al., 2017). Steroid-induced elevated IOP is due to the increased resistance in aqueous outflow pathway in the TM. Alterations in the function of TM can eventually lead to increased outflow resistance and elevated IOP (Clark and Wordinger, 2009). However, the molecular mechanism responsible for the pathogenesis of SI-OHT/SIG is poorly understood (Fini et al., 2017; Liesenborghs et al., 2020). Therefore, in the present study, the role of miRNAs in mediating differential GC responsiveness in HTM cells was investigated.

Multiple studies support a significant role for miRNAs in glaucoma pathogenesis and potential miRNAs were identified in glaucoma-affected tissues and fluids such as aqueous humor, tears, TM cells and retina which were derived from patients and animal models (Greene et al., 2022). In the trabecular meshwork, key miRNAs have been identified in TM cells from rodent and human subjected to cyclic mechanical stress, ROS or senescence (Li et al., 2010, 2009; Zhao et al., 2019; Shen et al., 2020; Youngblood et al., 2020). Identified miRNAs were found to have a crucial role in various cellular processes such as autophagy, apoptosis, senescence and neuro-inflammation pathways (Li et al., 2010, 2009; Zhao et al., 2019; Shen et al., 2020; Youngblood et al., 2020).

In TM cells, steroids are known to induce alterations in its structure and function including inhibition of cell proliferation and migration (Clark et al., 1994), cytoskeletal rearrangement (formation of cross-linked actins (CLANs) (Clark et al., 1995), increased TM cell and nuclear size (Wordinger et al., 1999), accumulation of excessive extracellular matrix (Zhou et al., 1998; Tane et al., 2007), decreased phagocytosis (Zhang et al., 2007) and alterations in cellular junctional complexes (Stamer and Clark, 2017). These cellular, biochemical and morphological changes ultimately lead to increased outflow resistance and hence elevated IOP. However, the expression of miRNAs in the TM in response to steroids has not yet been reported. Therefore, in the present study, the expression of miRNAs in GC-R and GC-NR HTM cells upon DEX treatment was investigated using small RNA sequencing technology.

Our study identified a unique miRNA signature between GC-R and GC-NR HTM cells. Specifically, 6 miRNAs were differentially expressed in GC-R HTM cells and 14 miRNAs were differentially expressed in GC-NR HTM cells. Two of these miRNAs associated with GC-NR (miR486-3p and miR-320a) were previously identified in aqueous humor of glaucoma patients. In the aqueous humor of glaucoma patients’ miR-486-3p was up-regulated but found to be down-regulated in GC-NR HTM cells (Tanaka et al., 2014). Conversely, miR-320a which was down-regulated in the aqueous humor of POAG patients was up-regulated in GC-NR HTM cells in the present study (Drewry et al., 2018). A number of DEMIRs have been identified in HTMcells in response to various stressors such as senescence, cyclic mechanical stress and oxidative stress; Li et al., 2010, 2009; Zhao et al., 2019; Shen et al., 2020; Youngblood et al., 2020). Only one of these DEMIRs was identified in the current study (miR-483-3p) and was a common miRNA between GC-R and GC-NR HTM cells (Shen et al., 2015). Mir-483-3p is known to affect the Wnt/β-catenin, the TGF-β, and the TP53 signaling pathways by targeting several genes as CTNNB1, SMAD4, IGF1, and BBC3 (Pepe et al., 2018). In HTM cells, the expression of miR-483-3p was reduced under H_2_O_2_-induced oxidative stress and the over-expression of 483-3p inhibited the expression of ECM proteins such as fibronectin, laminin and collagen by targeting TGFβ2/SMAD4 signaling (Shen et al., 2015). However, the role of miR-483-3p in steroid-induced OHT/glaucoma is not clear which warrants further investigation.

The unique DEMIR signature between GC-R and GC-NR HTM cells provides an opportunity to identify the molecular mechanisms driving GC-responsiveness and the development of SI-OHT and glaucoma. Uniquely miR-124-3p was significantly down-regulated in GC-R HTM cells as compared to GC-NR HTM cells. How this affects GC-responsiveness and alterations in the TM physiology is not known? but miR124-3p can negatively regulate the expression of GR by repressing the expression of the GC receptor (NR3C1) (Vreugdenhil et al., 2009; Wang et al., 2017). The expression of miR-124-3p and hence repression of NR3C1 is mediated via SMAD4 signaling; SMAD4 is a direct target of miR-483-3p (Liu et al., 2019; Shen et al., 2015). Elevated levels of miR-124-3p in the brain of rats and mice were associated with the decreased GR levels and GC sensitivity (Wang et al., 2017) and so potentially reduced miR-124-3p in GC-R HTM cells may suggest a role for miR-124-3p in the regulation of GR levels and GC sensitivity in the trabecular meshwork. Interestingly, miR-124-3p also modulates GR function indirectly by targeting phosphor-diesterase 4B or 11β-hydroxysteroid dehydrogenase 1(11β-HSD1) (Kim et al., 2015; Xu et al., 2017). 11β-HSD1 converts cortisone and cortisol and elevated levels of cortisol have been found in patients with POAG and the inhibitors of 11β-HSD1 reduced IOP in glaucoma patients (Rauz et al., 2003; Choi et al., 2019, 2017). However, the exact role of11β-HSD1in steroid-induced glaucoma is not completely understood. Further studies are warranted to understand the role of miR124-3p in in differential GC responsiveness and SI-OHT and glaucoma.

Our pathway analysis of the DEMIRs revealed several significant pathways found in both GC-R and GC-NR HTM cells (Supplementary Table S4a-S4e). When considering differential effects on pathway enrichment axon guidance signaling was significantly up-regulated pathway in GC-R (p=0.03) and significantly down-regulated (p=1.5E-05) in GC-NR HTM cells. Interestingly, GCs are shown to have a potential role in nervous system including stress response in neurons, synthesis of neurotransmitters, neuronal survival and differentiation (McEwen et al., 1986; Gould and Tanapat, 1999; Glick et al., 2000; Wang et al., 2013). However, the importance of axon guidance signaling pathway in HTM cells needs further investigation. The relaxin signaling was significantly down-regulated (p=0.03) in GC-R HTM cells and conversely up-regulated in GC-NR cells.

Relaxin is a polypeptide hormone produced by the corpus luteum and the decidua in females and by the prostate in males(Zloto et al., 2020). Traditionally this hormone was associated with parturition during pregnancy by relaxing collagen fibers in the pelvic region (Hisaw, 1926). Relaxin mediates its pharmacological action by activating a group of seven transmembrane G-protein coupled receptors (GPCRs): relaxin family peptide receptors. There are four relaxin receptors 1-4 (RXFP1-4). Human relaxin H1 and H2 can activate both the receptor isoforms RXFP1 and RXFP2 (Scott et al., 2012). RXFP3 is believed to function primarily in the mouse brain (Scott et al., 2005) while the RXFP4 stimulated appetite and activated colon motility to control food intake and glucose homeostasis in the mouse intestine (McGowan et al., 2008). Apart from its role in pregnancy, relaxin is produced in many tissues in mammals were it has vasodilatory and anti-fibrotic effects (Bathgate et al., 2021). In the eye, the IOP lowering property of relaxin was first demonstrated by Paterson et al.1963 following intra-mascular injection of relaxin in humans. RXFP1 is expressed in sections of the anterior segment of the eye where it has been localized to the uveal, corneoscleral and cribriform meshwork and Schlemm’s canal endothelium suggesting its role in regulating outflow facility and IOP homeostasis (Zloto et al., 2020). Furthermore, of relevance to TM changes in response to glucocorticoids, pre-clinical studies suggest the potential use of relaxin as anti-fibrotic molecule in several organs such as lung, skin, kidney, heart and liver (Chen et al., 2020). Given that relaxin signaling was down-regulated in GC-R HTM cells and relaxin mediates anti-fibrotic activity through its cognate receptor (RXFP1) (Ng et al., 2019), the role of relaxin in SI-OHT and glaucoma requires further studies.

Our integrative miRNA-mRNA analysis of both *in silico* and our own RNA sequencing data from paired TM donor cells with known GC responsiveness (Kathirvel et al., 2022) identified several potential miRNA-mRNA interactions highlighting GC –responsive and non-responsive molecular regulators. For example, in GC-R HTM cells the up-regulation of miR 2114-3p was associated with the down-regulation of several target genes such as NDP, GJB2, RAMP3, PDZK1P1, RTN4RL1, DRAXIN, KRT6, S100A14, CHRNA4, and RIC3. Whereas the down-regulation of miR-124-3p in the GC-R HTM cells was associated with the up-regulation of the transcription factor KLF-15. Interestingly in GC-NR group, miR-486-3p, miR-181b-1-3p and miR-10396b-5p were found to be molecular regulators in mediating GC-non-responsiveness by targeting several mRNA target genes.

In conclusion, this is the first study that identified a unique miRNA signature between GC-R and GC-NR HTM cells using small RNA sequencing. Ingenuity Pathway Analysis identified enriched pathways and biological processes associated with differential glucocorticoid (GC) responsiveness in HTM cells. Integrative analysis of miRNA-mRNA of the same set of HTM cells revealed several molecular regulators for GC non-responsiveness. Additional functional studies of these molecular regulators are warranted to validate their role in differential GC responsiveness. Understanding the role of miRNAs in GC responsiveness raises the possibility of new molecular targets for the management of steroid-OHT/glaucoma.

## Author Contributions

Conceptualization: SS; Methodology: SS, DB; Software: DB, KK; Validation: SS, DB; Formal Analysis: KK, RH, SS, DB; Investigation: KK, RH; Resources: SS, DB; Data Curation: SS, DB; Writing-Original Draft: KK, RH, SS, DB; Writing-Review and Editing: RK, VRM, RS, CEW; Supervision: SS, DB; Project administration: SS; Funding acquisition: SS.

## Institutional Review Board Statement

This study was approved by the standing Human Ethics Committee of Aravind Medical Research Foundation, Madurai, Tamilnadu, India (ID NO. RES2017006BAS).

## Informed Consent Statement

Not Applicable.

## Data Availability Statement

Data Access: The raw mRNA sequencing data of HTM cells from each human donor eye used in the present study have been deposited publicly in NCBI-SRA under the BioProject PRJNA982055 (https://www.ncbi.nlm.nih.gov/sra/PRJNA982055). Code Availability: The bioinformatics In-house pipeline used for mRNA sequencing data analysis in the present study have been submitted to GitHub in shell script (https://github.com/SenthilKumariLab/mRNA-seq-Analysis-Pipeline.git)

## Acknowledgments

The authors acknowledge the Rotary Aravind International Eye Bank, Aravind Eye Hospital, Madurai, India for providing human donor eyes for this study.

