## Supplementary Table for "Small RNA Sequencing Reveals a Distinct MicroRNA Signature between Glucocorticoid Responder and Glucocorticoid Non-responder Primary Human Trabecular Meshwork Cells after Dexamethasone Treatment"

**Table S1:** Characteristics of Human Trabecular Meshwork Cells Used for the Present Study

| **S.No** | **R/NR** | **Code** | **Age** | **Sex** | **Treatment**  **0.1%/100nM** |
| --- | --- | --- | --- | --- | --- |
| 1 | R | OCHD18-39 | 82 | F | ETH/DEX |
| 2 |  | OCHD18-53 | 67 | M | ETH/DEX |
| 3 |  | OCHD19-03 | 65 | F | ETH/DEX |
| 4 |  | OCHD19-04 | 72 | M | ETH/DEX |
| 5 | NR | OCHD18-49 | 48 | M | ETH/DEX |
| 6 |  | OCHD18-52 | 55 | F | ETH/DEX |
| 7 |  | OCHD18-56 | 82 | M | ETH/DEX |
| 8 |  | OCHD19-02 | 66 | M | ETH/DEX |

**Table S2:** Alignment Statistics of miRNA Sequencing Data

| **S. No** | **R/NR** | **ID** | | **Total Reads**  **(in millions)** | **% of Mapped Reads** | **% of**  **Unmapped**  **Reads** | **No. of miRNAs** |
| --- | --- | --- | --- | --- | --- | --- | --- |
| 1 | R | 18-39 | ETH | 17.2 | 98.2 | 1.8 | 892 |
|  |  |  | DEX | 14.6 | 98.2 | 1.8 | 898 |
| 2 |  | 18-53 | ETH | 11.3 | 97.3 | 2.7 | 828 |
|  |  |  | DEX | 12.1 | 97.4 | 2.6 | 841 |
| 3 |  | 19-03 | ETH | 13.2 | 98.3 | 1.7 | 871 |
|  |  |  | DEX | - | - | - | - |
| 4 |  | 19-04 | ETH | 13.8 | 98.4 | 1.6 | 856 |
|  |  |  | DEX | 14.2 | 98.5 | 1.5 | 857 |
| 5 | NR | 18-49 | ETH | 10.8 | 97.1 | 2.9 | 849 |
|  |  |  | DEX | 9.1 | 97.4 | 2.6 | 837 |
| 6 |  | 18-52 | ETH | 13.6 | 97.6 | 2.4 | 878 |
|  |  |  | DEX | 11.4 | 95.8 | 4.2 | 872 |
| 7 |  | 18-56 | ETH | 12.0 | 98.5 | 1.5 | 857 |
|  |  |  | DEX | 10.7 | 87.3 | 2.7 | 718 |
| 8 |  | 19-02 | ETH | 12.0 | 98.3 | 1.7 | 863 |
|  |  |  | DEX | 13.6 | 98.6 | 1.4 | 877 |
|  |  |  | **Mean** | **12** | **97** |  | **853** |

**Table S3a**. List of Up/Down-regulated miRNAs from Group #1

| **Up-regulated miRNAs** | | | |  | **Down-regulated miRNAs** | | | |
| --- | --- | --- | --- | --- | --- | --- | --- | --- |
| **miRNA** | **logFC** | **logCPM** | **P Value** |  | **miRNA** | **logFC** | **logCPM** | **P Value** |
| hsa-miR-2114-3p | 6.07 | -1.16 | 4.25E-04 |  | hsa-miR-335-5p | -1.7 | 6.8 | 0.02 |
| hsa-miR-675-3p | 4.86 | 0.11 | 8.01E-04 |  | hsa-miR-549a-5p | -2.1 | 0.9 | 0.02 |
| hsa-miR-483-3p | 4.61 | 2.88 | 1.01E-04 |  | hsa-miR-335-3p | -3.0 | 8.4 | 1.01E-04 |
| hsa-miR-675-5p | 4.59 | 1.20 | 2.88E-04 |  | hsa-miR-7151-3p | -3.1 | -0.3 | 0.03 |
| hsa-miR-483-5p | 4.31 | 0.82 | 1.16E-04 |  | hsa-miR-124-3p | -6.2 | -0.8 | 0.01 |
| hsa-miR-5690 | 3.61 | -1.26 | 0.00 |  |  |  |  |  |
| hsa-miR-2114-5p | 3.17 | -0.83 | 0.02 |  |  |  |  |  |
| hsa-miR-6842-3p | 1.50 | 1.33 | 0.02 |  |  |  |  |  |

Note: *Group A*: DE miRNAs between DEX and ETH treated GC-R HTM cells

**Table S3b**. List of Up/Down-regulated miRNAs from Group #2

| **Up-regulated miRNAs** | | | |  | |  | **Down-regulated miRNAs** | | | |
| --- | --- | --- | --- | --- | --- | --- | --- | --- | --- | --- |
| **miRNA** | **logFC** | **logCPM** | **P Value** |  | **miRNA** | | | **logFC** | **logCPM** | **P Value** |
| hsa-miR-4485-5p | 4.73 | 1.01 | 2.89E-04 |  | hsa-miR-181b-2-3p | | | -1.21 | 0.85 | 0.04 |
| hsa-miR-483-5p | 4.65 | 1.20 | 4.42E-06 |  | hsa-miR-486-3p | | | -1.34 | 2.38 | 0.05 |
| hsa-miR-12136 | 4.39 | 7.32 | 0.00 |  | hsa-miR-6853-3p | | | -1.80 | -0.69 | 0.03 |
| hsa-miR-4328 | 4.11 | -0.38 | 0.02 |  | hsa-miR-335-3p | | | -1.93 | 8.56 | 0.00 |
| hsa-miR-483-3p | 4.05 | 3.36 | 5.60E-05 |  | hsa-miR-550a-3p | | | -2.06 | -0.14 | 0.03 |
| hsa-miR-675-5p | 3.67 | 3.14 | 0.00 |  | hsa-miR-9-3p | | | -4.88 | -1.33 | 0.02 |
| hsa-miR-675-3p | 3.63 | 2.61 | 0.01 |  |  | | |  |  |  |
| hsa-miR-320a-5p | 3.47 | 1.78 | 0.00 |  |  | | |  |  |  |
| hsa-miR-4485-3p | 3.07 | 3.57 | 0.00 |  |  | | |  |  |  |
| hsa-miR-10396b-5p | 2.66 | -0.69 | 0.01 |  |  | | |  |  |  |
| hsa-miR-10396a-5p | 2.60 | -0.72 | 0.01 |  |  | | |  |  |  |
| hsa-miR-3195 | 2.44 | -0.73 | 0.04 |  |  | | |  |  |  |
| hsa-miR-1246 | 2.24 | 1.01 | 0.01 |  |  | | |  |  |  |
| hsa-miR-5690 | 2.15 | -0.79 | 0.02 |  |  | | |  |  |  |
| hsa-miR-6842-3p | 1.50 | 2.08 | 0.01 |  |  | | |  |  |  |

**Note:** *Group B:* DE miRNAs between DEX and ETH treated GC-NR HTM cells

**Table S4a.** List of Enriched Pathways from DE miRNAs of Group #1

| **Pathways associated with Up-Regulated miRNAs** | | | | |  | **Pathways associated with Down-Regulated miRNAs** | | | | |
| --- | --- | --- | --- | --- | --- | --- | --- | --- | --- | --- |
| **Pathways** | **PValue** | **Genes** | **Fold Enrichment** | **FDR** |  | **Pathways** | **PValue** | **Genes** | **Fold Enrichment** | **FDR** |
| Glioma | 6.94E-05 | PDGFRA, CAMK2D, PRKCA, EGFR, DDB2, PIK3CA, CDK4, MDM2, E2F2, MAPK1, PLCG1, CAMK2G, MAPK3 | 4.05 | 0.006 |  | Long-term depression | 3.09E-04 | GRIA1, MAP2K1, GUCY1A1, IGF1, GNAI2, GNA13, NRAS, PPP2R1A, GNAQ, GNA12, PLCB1, GRIA3, PRKG1 | 3.42 | 0.017 |
| Non-small cell lung cancer | 2.12E-04 | PIK3CA, PDPK1, CDK4, HGF, MAPK1, E2F2, PRKCA, PLCG1, STK4, EGFR, MAPK3, DDB2 | 3.89 | 0.012 |  | cGMP-PKG signaling pathway | 0.002 | MEF2A, MAP2K1, GUCY1A1, ROCK1, ATP2B4, NFATC1, ATP1A2, KNG1, GNAI2, GNA13, EDNRA, PPP3CB, AKT2, GNAQ, GNA12, AKT3, CREB3L2, PPIF, CALM1, PLCB1, PRKG1, SLC25A6 | 2.08 | 0.044 |
| Axon guidance | 7.40E-04 | SEMA7A, CAMK2D, BMPR2, SEMA4D, UNC5B, PDPK1, PRKCA, EFNA5, PPP3R1, PIK3CA, DPYSL5, PLXNA2, MAPK1, SLIT3, PLCG1, EPHB2, SRGAP2, CAMK2G, MAPK3 | 2.44 | 0.025 |  | Prostate cancer | 0.003 | GSK3B, MAP2K1, TMPRSS2, IGF1, CASP9, NRAS, AKT2, AKT3, CREB3L2, MDM2, BCL2, CTNNB1, GRB2, E2F3, FGFR2 | 2.44 | 0.047 |
| MAPK signaling pathway | 7.49E-04 | ARRB1, ARRB2, STK4, EFNA5, ELK1, EGFR, CACNG8, PPP3R1, MAPK8, FGF9, MAP3K20, MAPK1, MAPK3, PDGFRA, MAP4K2, NTRK2, MEF2C, HGF, PRKCA, CACNA2D4, DUSP8, MAPK8IP3, MAPK13, TRAF6, MAPKAPK2, MAP3K13 | 2.07 | 0.025 |  | PI3K-Akt signaling pathway | 0.003 | CHRM2, GSK3B, ITGB3, LAMC2, FGF1, EFNA5, CASP9, NRAS, BCL2L11, RXRA, ERBB3, GNG5, PPP2R1A, ERBB4, AKT2, AKT3, CREB3L2, IL6R, IFNAR2, NTRK2, MAP2K1, ANGPT1, ITGA3, BDNF, IGF1, PRLR, EREG, COL1A1, CDK6, RPS6KB1, DDIT4, ITGA11, GNB4, BCL2, MDM2, GRB2, FGFR2 | 1.65 | 0.047 |
| Focal adhesion | 9.25E-04 | PDGFRA, PDPK1, HGF, ITGB3, XIAP, PRKCA, ELK1, EGFR, ARHGAP35, MYLK, MAPK8, PIK3CA, TNN, COL6A1, RAPGEF1, PIP5K1A, MAPK1, PIP5K1B, PPP1R12B, MAPK3 | 2.33 | 0.025 |  | Growth hormone synthesis, secretion and action | 0.003 | MAP2K3, SHC4, STAT5A, GSK3B, MAP2K1, SHC1, IGFBP3, FOS, IGF1, GNAI2, NRAS, AKT2, GNAQ, AKT3, CREB3L2, GRB2, PLCB1 | 2.25 | 0.047 |
| Endocytosis | 0.001 | PDGFRA, RAB5B, RAB5C, IQSEC2, ASAP3, VPS37C, ARRB1, VPS26B, ARRB2, EGFR, ITCH, GRK3, GRK2, CAPZB, TRAF6, PSD4, PSD3, CHMP1A, MDM2, PIP5K1A, PIP5K1B, STAMBP, RAB11FIP4 | 2.14 | 0.026 |  | Estrogen signaling pathway | 0.006 | SHC4, MAP2K1, KCNJ6, SHC1, FOS, OPRM1, GNAI2, NRAS, SP1, AKT2, GNAQ, AKT3, CREB3L2, BCL2, GRB2, CALM1, PLCB1, KCNJ3 | 2.06 | 0.065 |
| FoxO signaling pathway | 0.001 | CDKN2B, SMAD4, PDPK1, SETD7, STK4, SOD2, PRKAB1, EGFR, GRM1, MAPK13, MAPK8, PIK3CA, MDM2, MAPK1, MAPK3 | 2.68 | 0.029 |  | Other types of O-glycan biosynthesis | 0.009 | GALNT7, POMT2, GALNT6, ST6GAL2, POFUT1, EOGT, GXYLT1, PLOD3, GALNT9 | 3.02 | 0.080 |
| Calcium signaling pathway | 0.003 | PDGFRA, NTRK2, CAMK2D, HGF, NTRK3, PTGER3, PRKCA, ATP2B1, EGFR, TPCN2, GRM1, MYLK, P2RX7, PPP3R1, GDNF, FGF9, ASPH, PHKG2, NOS1, PLCG1, CAMK2G | 2.04 | 0.054 |  | Signaling pathways regulating pluripotency of stem cells | 0.009 | GSK3B, MAP2K1, PCGF5, LIF, LIFR, IGF1, SMAD5, REST, NRAS, ACVR1C, AKT2, KAT6A, AKT3, CTNNB1, GRB2, BMPR1B, SKIL, FGFR2 | 1.99 | 0.080 |
| ErbB signaling pathway | 0.003 | CAMK2D, MAPK8, PIK3CA, MAPK1, PRKCA, NRG1, PLCG1, CAMK2G, ELK1, EGFR, MAPK3 | 3.02 | 0.054 |  | Bacterial invasion of epithelial cells | 0.009 | ACTR3, SHC4, SHC1, SEPTIN2, RHOG, CTNNB1, ELMO2, ARPC5, SEPTIN9, WASF2, SEPTIN11, CD2AP | 2.46 | 0.080 |
| Ras signaling pathway | 0.005 | PDGFRA, NTRK2, RAB5B, RALBP1, RAB5C, HGF, PRKCA, KSR2, STK4, EFNA5, ELK1, GRIN2B, EGFR, MAPK8, PIK3CA, FGF9, MAPK1, PLCG1, BCL2L1, MAPK3 | 1.99 | 0.076 |  | Apoptosis | 0.025 | DFFB, MAP2K1, CFLAR, FOS, CTSS, ERN1, CASP9, NRAS, BCL2L11, CASP10, AKT2, AKT3, BCL2, CTSH, CYCS, CTSB | 1.86 | 0.155 |
| GnRH secretion | 0.006 | KCNJ5, PIK3CA, MAPK1, ARRB1, PRKCA, KCNN3, ARRB2, KCNJ3, MAPK3 | 3.29 | 0.076 |  | Gastric cancer | 0.026 | SHC4, GSK3B, MAP2K1, SHC1, FGF1, TGFBR1, LRP6, NRAS, RXRA, RPS6KB1, AKT2, AKT3, BCL2, CTNNB1, GRB2, E2F3, FGFR2 | 1.80 | 0.159 |
| Rap1 signaling pathway | 0.008 | PDGFRA, HGF, ITGB3, CTNND1, FPR1, PRKCA, EFNA5, GRIN2B, EGFR, ADCY6, MAPK13, PIK3CA, FGF9, RAPGEF1, PFN4, MAPK1, PLCG1, MAPK3 | 2.00 | 0.102 |  | Renal cell carcinoma | 0.029 | RAP1B, MAP2K1, NRAS, AKT2, AKT3, RAPGEF1, GRB2, VHL, ELOC, PAK3 | 2.29 | 0.165 |
| VEGF signaling pathway | 0.012 | PPP3R1, PIK3CA, MAPKAPK2, MAPK1, PRKCA, PLCG1, MAPK13, MAPK3 | 3.17 | 0.126 |  | TNF signaling pathway | 0.050 | MAP2K3, MAP2K1, LIF, CFLAR, FOS, RPS6KA4, RPS6KA5, CASP10, TRAF3, CCL5, AKT2, AKT3, CREB3L2 | 1.83 | 0.239 |
| GnRH signaling pathway | 0.017 | CAMK2D, MAPK8, MAPK1, PRKCA, CAMK2G, ELK1, ADCY6, EGFR, MAPK13, MAPK3 | 2.51 | 0.163 |  | JAK-STAT signaling pathway | 0.050 | STAT5A, IFNAR2, IL11, LIF, LIFR, IFNLR1, TYK2, PRLR, PIAS2, PIAS1, AKT2, AKT3, BCL2, GRB2, IL6R, PTPN2, SOCS5 | 1.66 | 0.239 |
| Long-term potentiation | 0.023 | CAMK2D, PPP3R1, MAPK1, PRKCA, CAMK2G, GRIN2B, GRM1, MAPK3 | 2.79 | 0.206 |  | Insulin signaling pathway | 0.051 | SHC4, GSK3B, MAP2K1, SHC1, ACACA, NRAS, RPS6KB1, PPP1R3B, AKT2, AKT3, RAPGEF1, TRIP10, FLOT2, GRB2, CALM1 | 1.73 | 0.239 |
| Insulin secretion | 0.030 | CAMK2D, ADCYAP1R1, KCNMB2, KCNMA1, ATP1B4, PRKCA, KCNN3, CAMK2G, ADCY6 | 2.45 | 0.226 |  | SNARE interactions in vesicular transport | 0.054 | VAMP8, VAMP7, STX17, STX7, SNAP23, GOSR1 | 2.87 | 0.249 |
| T cell receptor signaling pathway | 0.033 | PPP3R1, MAPK8, PIK3CA, PDPK1, CDK4, MAPK1, PLCG1, ICOS, MAPK13, MAPK3 | 2.25 | 0.239 |  |  |  |  |  |  |
| Endometrial cancer | 0.036 | PIK3CA, PDPK1, MAPK1, ELK1, EGFR, MAPK3, DDB2 | 2.82 | 0.239 |  |  |  |  |  |  |
| Cell cycle | 0.043 | ORC4, CDKN2B, SMAD4, ANAPC15, ANAPC16, CDK4, MDM2, E2F2, CDC6, SMC1A, CDC25C | 2.04 | 0.247 |  |  |  |  |  |  |
| Regulation of actin cytoskeleton | 0.045 | PDGFRA, CYFIP1, ITGB3, IQGAP1, EGFR, ARHGAP35, MYLK, PIK3CA, FGF9, SPATA13, PIP5K1A, PFN4, MAPK1, PIP5K1B, PPP1R12B, MAPK3 | 1.72 | 0.259 |  |  |  |  |  |  |
| Relaxin signaling pathway | 0.049 | MAPK8, PIK3CA, MAPK1, ARRB1, PRKCA, ARRB2, NOS1, ADCY6, EGFR, MAPK13, MAPK3 | 1.99 | 0.271 |  |  |  |  |  |  |
| NOD-like receptor signaling pathway | 0.051 | XIAP, MAPK13, P2RX7, MAVS, MAPK8, DHX33, MAP1LC3C, OAS2, TRAF6, MFN2, MAPK1, TRPM7, BCL2L1, MAPK3 | 1.78 | 0.274 |  |  |  |  |  |  |

**Table S4b.** List of Enriched Pathways from DE miRNAs of Group #2

| **Pathways associated with Up-Regulated miRNAs** | | | | |  | **Pathways associated with Down-Regulated miRNAs** | | | | |
| --- | --- | --- | --- | --- | --- | --- | --- | --- | --- | --- |
| **Pathways** | **PValue** | **Genes** | **Fold Enrichment** | **FDR** |  | **Pathways** | **PValue** | **Genes** | **Fold Enrichment** | **FDR** |
| MAPK signaling pathway | 9.29E-05 | ARRB1, ARRB2, FGF1, ELK1, RAP1B, MAPK9, CACNG8, NRAS, PPP3R1, MAPK8, FGF9, MKNK2, MAP3K20, MAPK1, MAPK3, PDGFRA, MAP4K2, NTRK2, MEF2C, DUSP3, HGF, CACNA2D1, INSR, IGF2, PRKCA, DUSP8, MAPK8IP3, PPM1A, PPP5C, TAOK1, TRAF6, KIT, MAPKAPK2, RAPGEF2, MAPT, MAP3K13 | 2.009 | 0.007 |  | TGF-beta signaling pathway | 0.001 | GREM1, TGFB1, RBL1, PPP2R1A, NOG, BMP8A, MAPK1, HAMP, SMAD5, BMP7, ACVR2A, MAPK3 | 3.214 | 0.097 |
| Insulin secretion | 1.84E-04 | CAMK2D, ATP1B4, PRKCA, ADCY1, ATP1B1, ADCY7, ADCY6, CREB1, CCKAR, KCNMB2, KCNMA1, KCNN3, CAMK2G, STX1A, VAMP2, CREB5 | 3.053 | 0.008 |  | Regulation of actin cytoskeleton | 0.006 | ITGB1, NCKAP1, SRC, ITGA2, PXN, LIMK1, BRAF, ARPC4, FGF2, PTK2, KNG1, APC, MAPK1, NCKAP1L, ITGA5, PAK3, WASF2, MAPK3 | 2.078 | 0.163 |
| Long-term potentiation | 6.23E-04 | CAMK2D, PRKCA, ADCY1, GRIN2B, GRM1, RAP1B, GRIN2A, NRAS, PPP3R1, CAMK4, MAPK1, CAMK2G, MAPK3 | 3.184 | 0.015 |  | TNF signaling pathway | 0.013 | RPS6KA5, TRAF3, CCL5, CREB3L2, MAPK1, TRAF1, CFLAR, MAP3K14, CREB5, MAPK13, MAPK3 | 2.472 | 0.239 |
| Aldosterone synthesis and secretion | 7.85E-04 | KCNJ5, CAMK2D, NPR1, ATP2B3, ATP1B4, PRKCA, ATP2B1, ADCY1, ATP1B1, ADCY7, ADCY6, CREB1, CYP21A2, CAMK4, CAMK2G, CREB5 | 2.679 | 0.016 |  | p53 signaling pathway | 0.025 | CCND3, CDKN1A, TP53I3, MDM2, CDK1, IGF1, BCL2L1, RPRM | 2.759 | 0.287 |
| cAMP signaling pathway | 8.75E-04 | CAMK2D, NPR1, HHIP, PTGER3, ADCY1, ADCY7, ADCY6, RAP1B, MAPK9, GRIN2A, MAPK8, ADORA1, MAPK1, CAMK2G, MAPK3, ABCC4, PDE4D, ATP1B4, ATP2B3, ATP2B1, ATP1B1, GRIN2B, TSHR, CREB1, CAMK4, PDE3A, CREB5 | 2.005 | 0.017 |  | Signaling pathways regulating pluripotency of stem cells | 0.026 | ZFHX3, APC, PCGF5, DVL1, MAPK1, IGF1, FGF2, SMAD5, ACVR2A, MAPK13, WNT4, MAPK3 | 2.112 | 0.287 |
| Glioma | 0.002 | PDGFRA, CAMK2D, CDKN2A, PRKCA, DDB2, NRAS, CDK4, CAMK4, MDM2, MAPK1, PLCG1, CAMK2G, MAPK3 | 2.844 | 0.030 |  | Prostate cancer | 0.038 | CDKN1A, CREB3L2, MDM2, MAPK1, E2F2, BRAF, IGF1, CREB5, MAPK3 | 2.336 | 0.326 |
| Ras signaling pathway | 0.002 | RAB5B, RAB5C, FGF1, ELK1, RAP1B, MAPK9, GRIN2A, NRAS, MAPK8, GNG2, FGF9, RASSF5, MAPK1, PLCG1, PAK3, MAPK3, PDGFRA, NTRK2, HGF, INSR, IGF2, PRKCA, KSR2, GRIN2B, RALGAPB, KIT, BCL2L1 | 1.885 | 0.033 |  | PI3K-Akt signaling pathway | 0.041 | ITGB1, NTRK2, CDKN1A, ITGA2, TSC1, IGF1, EFNA5, FGF2, PRLR, PTK2, CCND3, PPP2R1A, ERBB4, CREB3L2, MDM2, GNB4, MAPK1, ITGA5, TLR4, CREB5, BCL2L1, MAPK3 | 1.564 | 0.332 |
| Focal adhesion | 0.002 | PDGFRA, PDPK1, CAV2, HGF, ITGB3, ITGA2, ITGA1, XIAP, PRKCA, PARVA, ELK1, ARHGAP35, RAP1B, MAPK9, MAPK8, TNN, ITGA8, RAPGEF1, PIP5K1A, MAPK1, PIP5K1B, PIP5K1C, PAK3, MAPK3 | 1.959 | 0.036 |  | Cell adhesion molecules | 0.047 | SPN, ITGB1, NLGN4Y, CNTNAP1, NFASC, OCLN, ALCAM, CADM1, SDC3, CNTN1, HLA-DPB1, CD276 | 1.924 | 0.332 |
| Axon guidance | 0.003 | CAMK2D, BMPR2, SEMA6A, SEMA4D, UNC5B, PDPK1, NFATC2, PRKCA, RND1, ENAH, ABLIM1, NRAS, PPP3R1, DPYSL5, PLXNA2, MAPK1, SLIT3, PLCG1, PAK3, CAMK2G, EPHB1, MAPK3 | 1.984 | 0.044 |  | C-type lectin receptor signaling pathway | 0.054 | CYLD, CD209, SRC, MDM2, MAPK1, CALM1, MAP3K14, MAPK13, MAPK3 | 2.178 | 0.361 |
| Non-small cell lung cancer | 0.004 | STAT5B, NRAS, PDPK1, CDKN2A, CDK4, RASSF5, HGF, MAPK1, PRKCA, PLCG1, MAPK3, DDB2 | 2.735 | 0.045 |  |  |  |  |  |  |
| Rap1 signaling pathway | 0.004 | PDGFRA, HGF, ITGB3, INSR, FPR1, PRKCA, ADCY1, FGF1, ADCY7, GRIN2B, ADCY6, RAP1B, ENAH, GRIN2A, NRAS, FGF9, RASSF5, KIT, RAPGEF1, MAPK1, RAPGEF2, LCP2, PLCG1, MAPK3 | 1.875 | 0.047 |  |  |  |  |  |  |
| ErbB signaling pathway | 0.005 | STAT5B, CAMK2D, PRKCA, NRG1, ELK1, MAPK9, NRAS, MAPK8, MAPK1, PLCG1, PAK3, CAMK2G, MAPK3 | 2.510 | 0.050 |  |  |  |  |  |  |
| Pathways in cancer | 0.005 | CAMK2D, CTBP2, EPO, HHIP, PTGER3, XIAP, ADCY1, FGF1, ELK1, ADCY7, ADCY6, BBC3, GNA13, MAPK9, NRAS, MAPK8, GNG2, FGF9, RASSF5, SUFU, MAPK1, PLCG1, CAMK2G, MAPK3, RUNX1T1, SMAD2, PDGFRA, STAT5B, CDKN2B, SMAD4, TPM3, CDKN2A, HGF, ITGA2, IGF2, PRKCA, MITF, DDB2, CCNA2, HEYL, AR, TRAF3, CDK4, TRAF6, KIT, RARA, MDM2, BCL2L1 | 1.483 | 0.050 |  |  |  |  |  |  |
| T cell receptor signaling pathway | 0.010 | PDPK1, NFATC2, MAPK9, NRAS, PPP3R1, MAPK8, CDK4, CD28, MAPK1, LCP2, PLCG1, PAK3, ICOS, MAPK3 | 2.209 | 0.089 |  |  |  |  |  |  |
| Relaxin signaling pathway | 0.011 | SMAD2, PRKCA, ARRB1, ADCY1, ARRB2, ADCY7, ADCY6, MAPK9, NRAS, MAPK8, CREB1, GNG2, MAPK1, NOS1, CREB5, MAPK3 | 2.035 | 0.097 |  |  |  |  |  |  |
| GnRH secretion | 0.015 | KCNJ5, NRAS, MAPK1, ARRB1, PRKCA, KCNN3, ARRB2, KCNJ3, HCN1, MAPK3 | 2.564 | 0.118 |  |  |  |  |  |  |
| Gap junction | 0.017 | PDGFRA, GUCY1A1, NRAS, GUCY1B1, MAPK1, PRKCA, ADCY1, ADCY7, TUBB4A, ADCY6, GRM1, MAPK3 | 2.238 | 0.131 |  |  |  |  |  |  |
| Insulin signaling pathway | 0.019 | SREBF1, PKLR, PDPK1, INSR, PHKB, ELK1, PRKAB1, MAPK9, NRAS, MAPK8, PRKAR1A, PHKG2, MKNK2, RAPGEF1, MAPK1, MAPK3 | 1.917 | 0.136 |  |  |  |  |  |  |
| Renin secretion | 0.023 | GUCY1A1, PPP3R1, GUCY1B1, CREB1, NPR1, KCNMA1, PDE3A, ADORA1, ADCY6, KCNJ2 | 2.378 | 0.152 |  |  |  |  |  |  |
| GnRH signaling pathway | 0.025 | MAPK9, CAMK2D, NRAS, MAPK8, MAPK1, PRKCA, ADCY1, ADCY7, CAMK2G, ELK1, ADCY6, MAPK3 | 2.117 | 0.158 |  |  |  |  |  |  |
| Transcriptional misregulation in cancer | 0.025 | MEF2C, DDX5, CCNT1, KMT2A, BCL11B, HMGA2, MITF, PBX1, DDB2, CDK9, HOXA10, CCNA2, NR4A3, MAF, SIX4, MDM2, RARA, H3-3B, BCL2L1, RUNX1T1 | 1.701 | 0.158 |  |  |  |  |  |  |
| Cellular senescence | 0.027 | SMAD2, CDKN2B, CDKN2A, FBXW11, NFATC2, HIPK1, HIPK3, CCNA2, NRAS, PPP3R1, CDK4, RASSF5, MAPKAPK2, MDM2, MAPK1, TRPM7, MAPK3 | 1.788 | 0.158 |  |  |  |  |  |  |
| FoxO signaling pathway | 0.028 | CDKN2B, SMAD4, PDPK1, INSR, SETD7, SOD2, FBXO32, PRKAB1, GRM1, MAPK9, NRAS, MAPK8, MDM2, MAPK1, MAPK3 | 1.879 | 0.158 |  |  |  |  |  |  |
| Oocyte meiosis | 0.028 | CAMK2D, ANAPC15, ANAPC16, FBXW11, ADCY1, CDC25C, SMC1A, YWHAZ, ADCY7, ADCY6, AR, PPP3R1, MAPK1, CAMK2G, MAPK3 | 1.879 | 0.158 |  |  |  |  |  |  |
| Long-term depression | 0.028 | GNA13, GUCY1A1, NRAS, GUCY1B1, MAPK1, PRKCA, NOS1, GRM1, MAPK3 | 2.462 | 0.158 |  |  |  |  |  |  |
| Growth hormone synthesis, secretion and action | 0.028 | STAT5B, PRKCA, ADCY1, SSTR3, ADCY7, ADCY6, MAPK9, NRAS, MAPK8, CREB1, MAPK1, PLCG1, CREB5, MAPK3 | 1.931 | 0.158 |  |  |  |  |  |  |
| Cell cycle | 0.042 | SMAD2, CDKN2B, SMAD4, ANAPC15, ANAPC16, CDKN2A, CDC6, CDC25C, SMC1A, YWHAZ, CCNA2, ORC4, CDK4, MDM2 | 1.823 | 0.201 |  |  |  |  |  |  |
| Pancreatic secretion | 0.045 | RAP1B, CCKAR, KCNMA1, ATP2B3, ATP1B4, PRKCA, ATP2B1, ADCY1, ATP1B1, ADCY7, ADCY6, TPCN2 | 1.931 | 0.207 |  |  |  |  |  |  |
| Regulation of lipolysis in adipocytes | 0.052 | NPR1, INSR, PTGER3, ADORA1, ADCY1, ADCY7, ADCY6, TSHR | 2.344 | 0.222 |  |  |  |  |  |  |
| Th1 and Th2 cell differentiation | 0.052 | MAPK9, STAT5B, PPP3R1, MAPK8, MAF, NFATC2, MAPK1, PLCG1, MAML3, RUNX3, MAPK3 | 1.962 | 0.222 |  |  |  |  |  |  |

**Table S4c.** List of Enriched Pathways from DE miRNAs of Group #3

| **Pathways associated with Up-Regulated miRNAs** | | | | |  | **Pathways associated with Down-Regulated miRNAs** | | | | |
| --- | --- | --- | --- | --- | --- | --- | --- | --- | --- | --- |
| **Pathways** | **PValue** | **Genes** | **Fold Enrichment** | **FDR** |  | **Pathways** | **PValue** | **Genes** | **Fold Enrichment** | **FDR** |
| Neurotrophin signaling pathway | 3.05E-04 | NTRK2, CAMK2D, PDPK1, NTRK3, MAPK8, ARHGDIA, TRAF6, MAPKAPK2, RAPGEF1, MAPK1, PLCG1, CAMK2G, MAPK3 | 3.508 | 0.031 |  | Adrenergic signaling in cardiomyocytes | 0.040 | CACNG8, RPS6KA5, PPP2R1A, ATP1A2, CALM1 | 3.829 | 1 |
| Glioma | 4.84E-04 | PDGFRA, CAMK2D, CDK4, MDM2, MAPK1, PRKCA, PLCG1, CAMK2G, MAPK3, DDB2 | 4.281 | 0.031 |  |  |  |  |  |  |
| Endocytosis | 7.81E-04 | PDGFRA, RAB5B, RAB5C, IQSEC2, ASAP3, VPS37C, ARRB1, ARRB2, GRK3, GRK2, CAPZB, TRAF6, PSD4, PSD3, CHMP1A, MDM2, PIP5K1A, PIP5K1B, STAMBP | 2.431 | 0.031 |  |  |  |  |  |  |
| Choline metabolism in cancer | 8.77E-04 | PDGFRA, MAPK8, SLC44A1, PDPK1, PIP5K1A, MAPK1, PRKCA, PIP5K1B, PLCG1, DGKK, MAPK3 | 3.604 | 0.031 |  |  |  |  |  |  |
| Oxytocin signaling pathway | 9.65E-04 | KCNJ5, MEF2C, CAMK2D, GUCY1B1, NPR1, PRKCA, ELK1, PRKAB1, ADCY6, PPP3R1, MAPK1, CAMK2G, KCNJ3, MAPK3 | 2.919 | 0.031 |  |  |  |  |  |  |
| MAPK signaling pathway | 0.002 | PDGFRA, MAP4K2, NTRK2, MEF2C, HGF, PRKCA, ARRB1, ARRB2, DUSP8, ELK1, MAPK8IP3, PPP3R1, MAPK8, FGF9, TRAF6, MAPKAPK2, MAP3K20, MAPK1, MAP3K13, MAPK3 | 2.184 | 0.045 |  |  |  |  |  |  |
| GnRH secretion | 0.003 | KCNJ5, MAPK1, ARRB1, PRKCA, KCNN3, ARRB2, KCNJ3, MAPK3 | 4.014 | 0.074 |  |  |  |  |  |  |
| Focal adhesion | 0.004 | PDGFRA, PDPK1, HGF, ITGB3, XIAP, PRKCA, ELK1, ARHGAP35, MAPK8, TNN, RAPGEF1, PIP5K1A, MAPK1, PIP5K1B, MAPK3 | 2.396 | 0.075 |  |  |  |  |  |  |
| Long-term potentiation | 0.005 | CAMK2D, PPP3R1, MAPK1, PRKCA, CAMK2G, GRIN2B, GRM1, MAPK3 | 3.834 | 0.075 |  |  |  |  |  |  |
| ErbB signaling pathway | 0.005 | CAMK2D, MAPK8, MAPK1, PRKCA, NRG1, PLCG1, CAMK2G, ELK1, MAPK3 | 3.400 | 0.075 |  |  |  |  |  |  |
| Non-small cell lung cancer | 0.007 | PDPK1, CDK4, HGF, MAPK1, PRKCA, PLCG1, MAPK3, DDB2 | 3.568 | 0.095 |  |  |  |  |  |  |
| FoxO signaling pathway | 0.007 | CDKN2B, SMAD4, MAPK8, PDPK1, SETD7, MDM2, MAPK1, SOD2, PRKAB1, GRM1, MAPK3 | 2.696 | 0.099 |  |  |  |  |  |  |
| Axon guidance | 0.011 | CAMK2D, BMPR2, SEMA4D, UNC5B, PDPK1, PRKCA, PPP3R1, DPYSL5, PLXNA2, MAPK1, PLCG1, CAMK2G, MAPK3 | 2.294 | 0.127 |  |  |  |  |  |  |
| EGFR tyrosine kinase inhibitor resistance | 0.011 | PDGFRA, HGF, MAPK1, PRKCA, NRG1, PLCG1, BCL2L1, MAPK3 | 3.252 | 0.127 |  |  |  |  |  |  |
| Ras signaling pathway | 0.014 | PDGFRA, NTRK2, RAB5B, RAB5C, HGF, PRKCA, KSR2, ELK1, GRIN2B, MAPK8, FGF9, MAPK1, PLCG1, BCL2L1, MAPK3 | 2.050 | 0.153 |  |  |  |  |  |  |
| Cell cycle | 0.016 | ORC4, CDKN2B, SMAD4, ANAPC15, ANAPC16, CDK4, MDM2, CDC6, SMC1A, CDC25C | 2.548 | 0.159 |  |  |  |  |  |  |
| Insulin secretion | 0.017 | CAMK2D, KCNMB2, KCNMA1, ATP1B4, PRKCA, KCNN3, CAMK2G, ADCY6 | 2.987 | 0.161 |  |  |  |  |  |  |
| GnRH signaling pathway | 0.025 | CAMK2D, MAPK8, MAPK1, PRKCA, CAMK2G, ELK1, ADCY6, MAPK3 | 2.762 | 0.199 |  |  |  |  |  |  |
| Pancreatic cancer | 0.030 | SMAD4, MAPK8, CDK4, MAPK1, BCL2L1, MAPK3, DDB2 | 2.958 | 0.213 |  |  |  |  |  |  |
| Aldosterone synthesis and secretion | 0.032 | KCNJ5, CAMK2D, NPR1, ATP1B4, PRKCA, ATP2B1, CAMK2G, ADCY6 | 2.621 | 0.215 |  |  |  |  |  |  |
| cGMP-PKG signaling pathway | 0.035 | MEF2C, PPP3R1, GUCY1B1, NPR1, KCNMB2, KCNMA1, MAPK1, ATP1B4, ATP2B1, ADCY6, MAPK3 | 2.115 | 0.223 |  |  |  |  |  |  |
| VEGF signaling pathway | 0.036 | PPP3R1, MAPKAPK2, MAPK1, PRKCA, PLCG1, MAPK3 | 3.265 | 0.223 |  |  |  |  |  |  |
| Long-term depression | 0.038 | GUCY1B1, MAPK1, PRKCA, NOS1, GRM1, MAPK3 | 3.211 | 0.229 |  |  |  |  |  |  |
| Pathways in cancer | 0.040 | CAMK2D, PTGER3, XIAP, ELK1, ADCY6, BBC3, MAPK8, FGF9, SUFU, MAPK1, PLCG1, CAMK2G, MAPK3, PDGFRA, CDKN2B, SMAD4, HGF, PRKCA, MITF, DDB2, HEYL, CDK4, TRAF6, MDM2, BCL2L1 | 1.512 | 0.235 |  |  |  |  |  |  |
| T cell receptor signaling pathway | 0.042 | PPP3R1, MAPK8, PDPK1, CDK4, MAPK1, PLCG1, ICOS, MAPK3 | 2.470 | 0.245 |  |  |  |  |  |  |
| Gap junction | 0.055 | PDGFRA, GUCY1B1, MAPK1, PRKCA, ADCY6, GRM1, MAPK3 | 2.554 | 0.305 |  |  |  |  |  |  |

**Table S4d.** List of Enriched Pathways from DE miRNAs of Group #4

| **Pathways associated with Up-Regulated miRNAs** | | | | |  | **Pathways associated with Down-Regulated miRNAs** | | | | |
| --- | --- | --- | --- | --- | --- | --- | --- | --- | --- | --- |
| **Pathways** | **PValue** | **Genes** | **Fold Enrichment** | **FDR** |  | **Pathways** | **PValue** | **Genes** | **Fold Enrichment** | **FDR** |
| Oxytocin signaling pathway | 0.004 | MEF2C, CACNG8, PPP3R1, EEF2K, CACNA2D4, PPP1R12B, EGFR, MYLK | 3.960 | 0.422 |  | EGFR tyrosine kinase inhibitor resistance | 4.08E-04 | SHC4, GSK3B, MAP2K1, SHC1, NRAS, BCL2L11, ERBB3, RPS6KB1, AKT2, AKT3, BCL2, GRB2, IL6R, FGFR2 | 3.149 | 0.024 |
| Axon guidance | 0.031 | SEMA7A, PPP3R1, PIK3CA, SLIT3, EPHB2, SRGAP2, EFNA5 | 2.932 | 1.000 |  | Long-term depression | 0.002 | GNA13, GRIA1, MAP2K1, GUCY1A1, NRAS, GNAQ, GNA12, PLCB1, PRKG1, GRIA3, GNAI2 | 3.258 | 0.052 |
| MAPK signaling pathway | 0.037 | NTRK2, MEF2C, CACNG8, PPP3R1, CACNA2D4, STK4, EFNA5, EGFR, MAPK13 | 2.333 | 1.000 |  | Growth hormone synthesis, secretion and action | 0.003 | MAP2K3, SHC4, STAT5A, GSK3B, MAP2K1, SHC1, IGFBP3, FOS, GNAI2, NRAS, AKT2, GNAQ, AKT3, CREB3L2, GRB2, PLCB1 | 2.389 | 0.065 |
|  |  |  |  |  |  | PI3K-Akt signaling pathway | 0.009 | CHRM2, GSK3B, ITGB3, LAMC2, FGF1, EFNA5, CASP9, NRAS, BCL2L11, RXRA, ERBB3, GNG5, AKT2, AKT3, CREB3L2, IL6R, IFNAR2, NTRK2, MAP2K1, ANGPT1, ITGA3, BDNF, PRLR, EREG, COL1A1, CDK6, RPS6KB1, DDIT4, ITGA11, BCL2, GRB2, FGFR2 | 1.606 | 0.087 |
|  |  |  |  |  |  | Gastric cancer | 0.009 | SHC4, GSK3B, MAP2K1, SHC1, FGF1, TGFBR1, LRP6, NRAS, RXRA, RPS6KB1, AKT2, AKT3, BCL2, CTNNB1, GRB2, E2F3, FGFR2 | 2.027 | 0.087 |
|  |  |  |  |  |  | Signaling pathways regulating pluripotency of stem cells | 0.014 | GSK3B, MAP2K1, LIF, LIFR, SMAD5, REST, NRAS, ACVR1C, AKT2, KAT6A, AKT3, CTNNB1, GRB2, BMPR1B, SKIL, FGFR2 | 1.988 | 0.111 |
|  |  |  |  |  |  | JAK-STAT signaling pathway | 0.019 | STAT5A, IFNAR2, IL11, LIF, LIFR, IFNLR1, TYK2, PRLR, PIAS2, PIAS1, AKT2, AKT3, BCL2, GRB2, IL6R, PTPN2, SOCS5 | 1.865 | 0.146 |
|  |  |  |  |  |  | Relaxin signaling pathway | 0.029 | SHC4, MAP2K1, SHC1, FOS, TGFBR1, GNAI2, COL1A1, NRAS, GNG5, AKT2, AKT3, CREB3L2, GRB2, PLCB1 | 1.928 | 0.184 |
|  |  |  |  |  |  | Lipid and atherosclerosis | 0.033 | MAP2K3, VAV3, GSK3B, NFATC1, VLDLR, FOS, SOD2, MIB1, ERN1, CASP9, PPP3CB, NRAS, RXRA, TRAF3, AKT2, AKT3, BCL2, CYCS, CD36, PLCB1 | 1.653 | 0.198 |
|  |  |  |  |  |  | Small cell lung cancer | 0.033 | CASP9, RXRA, CDK6, TRAF3, ITGA3, AKT2, AKT3, BCL2, CYCS, LAMC2, E2F3 | 2.125 | 0.198 |
|  |  |  |  |  |  | Focal adhesion | 0.033 | SHC4, VAV3, GSK3B, MAP2K1, ROCK1, SHC1, ITGA3, ITGB3, LAMC2, COL1A1, AKT2, AKT3, ITGA11, RAPGEF1, BCL2, CTNNB1, GRB2, PAK3, PPP1R12B | 1.680 | 0.198 |
|  |  |  |  |  |  | Neurotrophin signaling pathway | 0.035 | SHC4, NTRK2, GSK3B, MAP2K1, SHC1, BDNF, RPS6KA6, NRAS, AKT2, AKT3, RAPGEF1, BCL2, GRB2 | 1.941 | 0.198 |
|  |  |  |  |  |  | B cell receptor signaling pathway | 0.040 | VAV3, GSK3B, MAP2K1, PPP3CB, NRAS, AKT2, AKT3, NFATC1, GRB2, FOS | 2.167 | 0.218 |
|  |  |  |  |  |  | Insulin signaling pathway | 0.044 | SHC4, GSK3B, MAP2K1, SHC1, ACACA, NRAS, RPS6KB1, PPP1R3B, AKT2, AKT3, RAPGEF1, TRIP10, FLOT2, GRB2 | 1.816 | 0.231 |
|  |  |  |  |  |  | p53 signaling pathway | 0.051 | CASP9, STEAP3, CDK6, CDKN2A, ZMAT3, IGFBP3, CHEK1, BCL2, CYCS | 2.191 | 0.242 |

**Table S4e.** List of Enriched Pathways from DE miRNAs of Group #5

| **Pathways associated with Up-Regulated miRNAs** | | | | |  | **Pathways associated with Down-Regulated miRNAs** | | | | |
| --- | --- | --- | --- | --- | --- | --- | --- | --- | --- | --- |
| **Pathways** | **PValue** | **Genes** | **Fold Enrichment** | **FDR** |  | **Pathways** | **PValue** | **Genes** | **Fold Enrichment** | **FDR** |
| cGMP-PKG signaling pathway | 0.001 | GUCY1A1, INSR, NFATC2, ATP2B3, ADCY1, ATP1B1, ADCY7, ADRA1A, PIK3CG, PIK3R5, GNA13, CREB1, ADORA1, PDE3A, CREB5 | 2.703 | 0.118 |  | Axon guidance | 1.50E-05 | ITGB1, SEMA6A, SRC, LIMK1, SEMA4F, UNC5D, EFNA5, BMP7, PTK2, DPYSL5, ABL1, MAPK1, PLXNC1, SRGAP3, EPHB2, PAK3, SRGAP2, WNT4, MAPK3 | 3.313 | 0.004 |
| Insulin secretion | 0.002 | CCKAR, CREB1, PRKCA, KCNN3, ADCY1, ATP1B1, ADCY7, STX1A, VAMP2, CREB5 | 3.500 | 0.146 |  | Regulation of actin cytoskeleton | 0.009 | ITGB1, SRC, ITGA2, PXN, LIMK1, BRAF, ARPC4, FGF2, PTK2, APC, MAPK1, ITGA5, PAK3, WASF2, MAPK3 | 2.184 | 0.273 |
| MAPK signaling pathway | 0.004 | NTRK2, DUSP3, CACNA2D1, INSR, IGF2, PRKCA, FGF1, DUSP8, RAP1B, MAPK9, PPM1A, PPP5C, CACNG8, NRAS, TAOK1, KIT, MAPKAPK2, MKNK2, RAPGEF2, MAPT | 2.047 | 0.189 |  | TGF-beta signaling pathway | 0.009 | TGFB1, NOG, BMP8A, MAPK1, HAMP, SMAD5, BMP7, ACVR2A, MAPK3 | 3.038 | 0.273 |
| Oxytocin signaling pathway | 0.005 | GUCY1A1, CACNA2D1, NFATC2, PRKCA, ADCY1, ADCY7, PIK3CG, PIK3R5, CACNG8, NRAS, CAMK4, KCNJ2, KCNJ3 | 2.541 | 0.201 |  | ErbB signaling pathway | 0.017 | CDKN1A, SRC, ABL1, MAPK1, BRAF, PAK3, PTK2, MAPK3 | 2.987 | 0.365 |
| cAMP signaling pathway | 0.006 | ABCC4, HHIP, PDE4D, ATP2B3, ADCY1, ATP1B1, ADCY7, TSHR, RAP1B, MAPK9, GRIN2A, CREB1, CAMK4, ADORA1, PDE3A, CREB5 | 2.179 | 0.226 |  | Renal cell carcinoma | 0.021 | CDKN1A, TGFB1, EGLN2, MAPK1, BRAF, PAK3, MAPK3 | 3.220 | 0.365 |
| Aldosterone synthesis and secretion | 0.016 | CREB1, CYP21A2, CAMK4, ATP2B3, PRKCA, ADCY1, ATP1B1, ADCY7, CREB5 | 2.764 | 0.491 |  | Small cell lung cancer | 0.025 | ITGB1, CDKN1A, TRAF3, ITGA2, E2F2, TRAF1, PTK2, BCL2L1 | 2.760 | 0.365 |
| Ras signaling pathway | 0.024 | NTRK2, INSR, IGF2, PRKCA, KSR2, FGF1, RAP1B, MAPK9, GRIN2A, NRAS, GNG2, RALGAPB, RASSF5, KIT, PAK3 | 1.921 | 0.606 |  | Pancreatic cancer | 0.032 | CDKN1A, TGFB1, MAPK1, E2F2, BRAF, BCL2L1, MAPK3 | 2.923 | 0.365 |
| Salivary secretion | 0.032 | GUCY1A1, ATP2B3, PRKCA, ADCY1, ATP1B1, ADCY7, ADRA1A, VAMP2 | 2.617 | 0.606 |  | Signaling pathways regulating pluripotency of stem cells | 0.036 | ZFHX3, APC, DVL1, MAPK1, FGF2, SMAD5, ACVR2A, MAPK13, WNT4, MAPK3 | 2.219 | 0.365 |
| Growth hormone synthesis, secretion and action | 0.043 | MAPK9, STAT5B, NRAS, CREB1, PRKCA, ADCY1, ADCY7, SSTR3, CREB5 | 2.276 | 0.606 |  | VEGF signaling pathway | 0.037 | SRC, PXN, MAPK1, PTK2, MAPK13, MAPK3 | 3.227 | 0.365 |
| Rap1 signaling pathway | 0.045 | INSR, PRKCA, ADCY1, FGF1, ADCY7, RAP1B, ENAH, GRIN2A, NRAS, RASSF5, KIT, RAPGEF2, LCP2 | 1.863 | 0.606 |  | Long-term depression | 0.039 | GNAO1, GUCY1A1, MAPK1, BRAF, PRKG1, MAPK3 | 3.174 | 0.365 |

**Table S5**. List of miRNAs Selected from miRNA sequencing for Validation by qPCR

| **S.No** | **Symbol** |
| --- | --- |
| 1 | hsa-miR-124-3p |
| 2 | hsa-miR-335-3p |
| 3 | hsa-miR-335-5p |
| 4 | hsa-miR-483-3p |
| 5 | hsa-miR-483-5p |
| 6 | hsa-miR-549a-5p |
| 7 | hsa-miR-675-3p |
| 8 | hsa-miR-675-5p |
| 9 | hsa-miR-44853-3p |

**Table S6 showing the overlapping biological processes in ‘Target MRNA List 1 & 2’ from Group#3-#5**

| (A) *Biological processes in the ‘Target MRNA List 1 & 2’ from Group#3* | | | | |
| --- | --- | --- | --- | --- |
| Biological Process | **miRNA Target list 2** | | **miRNA Target list 1** | |
|  | **P-value** | **Adjusted P-value** | **P-value** | **Adjusted P-value** |
| Positive regulation of pri-miRNA transcription by RNA polymerase II (GO:1902895) | 0.005092 | 0.019881 | 0.006883 | 0.573186 |
| Regulation of pri-miRNA transcription by RNA polymerase II (GO:1902893) | 0.006735 | 0.019881 | 0.009904 | 0.618899 |
| (B) *Biological processes in the ‘Target MRNA List 1 & 2’ from Group#4* | | | | |
| Positive regulation of peptidyl-serine phosphorylation (GO:0033138) | 0.00246 | 0.055217 | 0.020185 | 0.479763 |
| Regulation of peptidyl-serine phosphorylation (GO:0033135) | 0.002727 | 0.055217 | 0.028546 | 0.479763 |
| Negative regulation of protein acetylation (GO:1901984) | 0.003994 | 0.055217 | 0.030568 | 0.479763 |
| amylin receptor signaling pathway (GO:0097647) | 0.004791 | 0.055217 | 0.044088 | 0.508983 |
| Glomerular visceral epithelial cell differentiation (GO:0072112) | 0.00956 | 0.055217 | 0.003997 | 0.313688 |
| regulation of axon regeneration (GO:0048679) | 0.00956 | 0.055217 | 0.029783 | 0.479763 |
| Positive regulation of calcium ion import (GO:0090280) | 0.010353 | 0.055217 | 0.037065 | 0.501795 |
| Neurotrophin TRK receptor signaling pathway (GO:0048011) | 0.011937 | 0.060559 | 0.009572 | 0.376974 |
| Regulation of granulocyte chemotaxis (GO:0071622) | 0.015886 | 0.064227 | 0.026916 | 0.479763 |
| Negative regulation of response to wounding (GO:1903035) | 0.016674 | 0.064227 | 0.031747 | 0.481647 |
| Neurotrophin signaling pathway (GO:0038179) | 0.016674 | 0.064227 | 0.031747 | 0.481647 |
| regulation of protein localization to cell surface (GO:2000008) | 0.026085 | 0.069851 | 0.011425 | 0.414312 |
| Central nervous system neuron differentiation (GO:0021953) | 0.026866 | 0.069851 | 0.013201 | 0.422548 |
| Positive regulation of protein phosphorylation (GO:0001934) | 0.034691 | 0.075955 | 0.030639 | 0.479763 |
| Receptor internalization (GO:0031623) | 0.038502 | 0.080893 | 0.007239 | 0.354107 |
| Plasma membrane bounded cell projection morphogenesis (GO:0120039) | 0.040813 | 0.082419 | 0.031381 | 0.481647 |
| Receptor metabolic process (GO:0043112) | 0.045421 | 0.085113 | 0.001852 | 0.269147 |
| (C) *Biological processes in the ‘Target MRNA List 1 & 2’ from Group#5* | | | | |
| Melanosome transport (GO:0032402) | 0.01164 | 0.051402 | 0.004006 | 0.229032 |
| Pigment granule transport (GO:0051904) | 0.01164 | 0.051402 | 0.004006 | 0.229032 |
| Regulation of brown fat cell differentiation (GO:0090335) | 0.01164 | 0.051402 | 0.004006 | 0.229032 |
| Establishment of melanosome localization (GO:0032401) | 0.012283 | 0.051402 | 0.005963 | 0.293283 |
| Long-term synaptic potentiation (GO:0060291) | 0.013568 | 0.051402 | 0.003091 | 0.20335 |
| Melanosome localization (GO:0032400) | 0.013568 | 0.051402 | 0.011939 | 0.382983 |
| NAD biosynthetic process (GO:0009435) | 0.013568 | 0.051402 | 0.011939 | 0.382983 |
| Peptidyl-threonine dephosphorylation (GO:0035970) | 0.01421 | 0.051402 | 0.01619 | 0.40656 |
| nucleotide metabolic process (GO:0009117) | 0.014852 | 0.051402 | 0.021433 | 0.456601 |
| Regulation of interleukin-4 production (GO:0032673) | 0.014852 | 0.051402 | 0.021433 | 0.456601 |
| Regulation of cytokine production involved in immune response (GO:0002718) | 0.015493 | 0.051402 | 0.002361 | 0.17261 |
| L-alpha-amino acid transmembrane transport (GO:1902475) | 0.017414 | 0.052623 | 0.006438 | 0.299846 |
| L-amino acid transport (GO:0015807) | 0.018053 | 0.052623 | 0.008608 | 0.331661 |
| Negative regulation of Notch signaling pathway (GO:0045746) | 0.019331 | 0.054048 | 0.001355 | 0.128933 |
| Positive regulation of cytokine production involved in immune response (GO:0002720) | 0.021245 | 0.05707 | 0.028641 | 0.475122 |
| Positive regulation of production of molecular mediator of immune response (GO:0002702) | 0.024427 | 0.059508 | 0.00143 | 0.129778 |
| Neurogenesis (GO:0022008) | 0.028234 | 0.059508 | 0.006668 | 0.305183 |
| Positive regulation of fat cell differentiation (GO:0045600) | 0.032657 | 0.065794 | 0.000576 | 0.081384 |
| Positive regulation of synaptic transmission (GO:0050806) | 0.046438 | 0.084827 | 0.000263 | 0.053254 |

*Table S6 shows the overlapping GO biological processes in the ‘Target MRNA List 1’ and ‘Target MRNA List 2’ from Group #3-#5 predicted by GSEApy package /python (v 3.8.3) (ranked by the smallest p-values of the biological processes in ‘Target MRNA List 2’). P-value<0.05 was considered to be statistically significant.*
